# Genomic Epidemiology with Mixed Samples

**DOI:** 10.1101/2020.04.03.021501

**Authors:** Tommi Mäklin, Teemu Kallonen, Jarno Alanko, Ørjan Samuelsen, Kristin Hegstad, Veli Mäkinen, Jukka Corander, Eva Heinz, Antti Honkela

**Affiliations:** Helsinki Institute for Information Technology HIIT, Department of Mathematics and Statistics, University of Helsinki, Helsinki, Finland; Department of Biostatistics, University of Oslo, Oslo, Norway; Wellcome Sanger Institute, Hinxton, Cambridgeshire, UK; Helsinki Institute for Information Technology HIIT, Department of Computer Science, University of Helsinki, Helsinki, Finland; Norwegian National Advisory Unit on Detection of Antimicrobial Resistance, Department of Microbiology and Infection Control, University Hospital of North Norway, Tromsø, Norway; Department of Pharmacy, UiT The Arctic University of Norway, Tromsø, Norway; Research group for Host-Microbe Interactions, Department of Medical Biology, Faculty of Health Sciences, UiT The Arctic University of Norway, Tromsø, Norway; Liverpool School of Tropical Medicine, Liverpool, United Kingdom

**Keywords:** genomic epidemiology, pathogen surveillance, plate sweeps, probabilistic modelling, pseudoalignment

## Abstract

Genomic epidemiology is a tool for tracing transmission of pathogens based on whole-genome sequencing. We introduce the mGEMS pipeline for genomic epidemiology with plate sweeps representing mixed samples of a target pathogen, skipping the colony pick step. The pipeline includes the novel mGEMS read binner for probabilistic assignments of sequencing reads, and the scalable pseudoaligner Themisto. We demonstrate the effectiveness of our approach using closely related samples in a nosocomial setting, obtaining results that are comparable to those based on colony picks. Our results lend firm support to more widespread consideration of genomic epidemiology with mixed infection samples.

## Background

Public health epidemiology for bacterial infections has been transformed by the use of high-throughput sequencing data to analyse and identify the source of an outbreak and to trace circulating pathogenic strains based on routine surveillance [1–5]. Standard genome-based epidemiological linking of cases requires accurate genome sequences for the pathogens derived from high coverage sequencing data for pure-colony isolates. The isolates are obtained by an enrichment and separation step in the form of a plate culture and subsequent colony picks based e.g. on morphology and colour. Typical workflow of genomic epidemiology may thus necessitate multiple colony picks per sample and the corresponding DNA library preparation and sequencing steps for each of them. Combined, these steps require a significant amount of laboratory effort and time, and lead to increased costs since the price of library preparation is becoming comparable to the cost of sequencing itself [6]. This can act as a barrier to more widespread genomic pathogen surveillance even in well-resourced public health laboratories, and prevent application of genomic epidemiology altogether in poorer settings.

Whole-genome shotgun metagenomics has been proposed as a solution for getting rid of the culturing step entirely. In this approach, sequencing is performed directly on the DNA extracted from the original sample and the resulting reads computationally binned or assembled. While tools capable of pangenome-based analyses [7], metagenome assembly [8–11], or taxonomic binning [12–14] from metagenomic short-read sequencing data have been developed, these methods typically require that the samples do not contain many closely related organisms. In particular, the strain-variation within a species is assumed to be large enough not to be confused with sequencing errors or variation in the assembly graph [15]. When more complex strain-level diversity is present, benchmarking these tools shows reduced performance in both taxonomic binning and metagenomic assembly [16–19]. In practice, natural strain-level variation is harbored ubiquitously in epidemiologically relevant samples [20–29] and it is reflected by the transmission events occurring between individuals and their environment [30]. Although some sample types may be dominated by one or two strains [31], direct sequencing of clinical samples may result in an overabundance of host DNA [28,32–34], or lack detection power for strains with low abundance in environments with high species diversity [18,32,35]. These challenges are overcome in genomic epidemiology by enriching the target species through the use of plate cultures. Since established protocols and growth media are available for most bacteria of clinical relevance [36], enrichment provides an effective means to deplete the host DNA and increase the sequencing depth for target organisms when working with well-characterized species.

In this article, we introduce the mGEMS pipeline for performing genomic epidemiology with mixed cultures from samples that may harbor multiple closely related bacterial lineages. mGEMS requires only a single culturing and library preparation step per sample, which can significantly reduce the cost of performing genomic epidemiology in the standard public health setting and make the whole process more streamlined. We demonstrate the effectiveness of our approach in SNP calling and phylogenetic analyses by using *in vitro* mixed samples of *Escherichia coli* and *Enterococcus faecalis* strains, as well as DNA reads synthetically mixed from closely related samples obtained from previous genomic epidemiology studies [29,37,38] tracking *E. faecalis*, *E. coli* and *Staphylococcus aureus* in public health settings. Likewise, the *E. coli* and *E. faecalis* strains used in the *in vitro* samples were hospital isolates and selected as representatives of clinically highly relevant sequence types. Our results illustrate that accurate transmission and case-linking analyses are possible at reduced cost levels by enabling sample de-mixing and subsequent variant calling.

Key parts of our pipeline presented in this paper are the mGEMS binner for short-read sequencing data, and the scalable pseudoaligner Themisto, which provides an exact version of the kallisto pseudoalignment algorithm [39] for large reference databases of single-clone sequenced bacterial pathogens. Together with recent advances in both probabilistic modelling of mixed bacterial samples [40] and genome assembly techniques [41], these methods form the mGEMS pipeline. A central step in mGEMS is an application of the recent mSWEEP method [40], which estimates the relative abundance of reference bacterial lineages in mixed samples using pseudoalignment and Bayesian mixture modelling. While Themisto enables upscaling of mSWEEP to significantly larger reference databases, the mGEMS binner is a novel sequencing read binning approach. Our binner is based on leveraging probabilistic sequencing read classifications to reference lineages from mSWEEP, and notably allowing a single read to be assigned to multiple bins. Using mGEMS to bin the reads in the original mixed samples produces sets of reads closely resembling standard isolate sequencing data and additionally acts as a denoising step for removing possible contaminant DNA. These advances allow a subsequent efficient use of the existing leading tools for genomic epidemiology in the analysis of mixed culture samples, which can pave way to a more widespread consideration of genomic epidemiology for public health applications.

## Results

### Read binning and genome assembly from mixed samples with mGEMS

Our mGEMS read binning algorithm, part of the mGEMS pipeline (Figure 1), requires probabilistic assignments of sequencing reads to reference taxonomic units (lineages or sequences) and an estimate of the relative sequence abundance of these same references in the full set of reads. mGEMS then bins the reads by assigning a read to a bin (corresponding to a target sequence from a given reference lineage) if the read-level assignment probability of the lineage is greater than or equal to the sequence abundance of that particular lineage in the full set of reads. Notably, this algorithm allows a single sequencing read to be assigned to multiple bins which is a crucial feature for considering strain-level variation. As shown in the Methods section, this algorithm assigns reads to reference lineages only if the sequence represented by a read is likely contained in a target sequence that belongs to the reference lineage.

**Figure 1.**
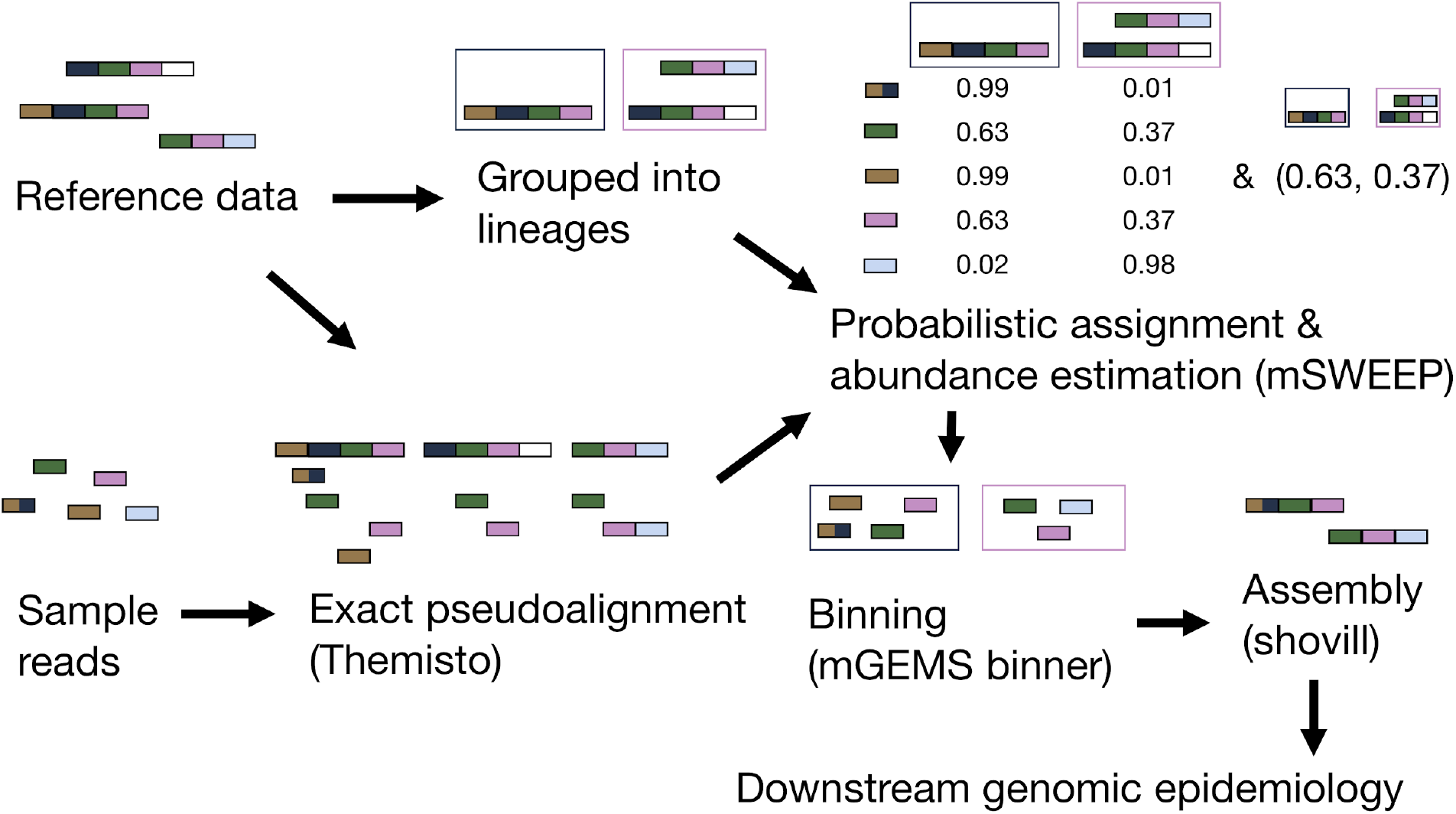
Flowchart describing a genomic epidemiology workflow with the mGEMS pipeline. The figure shows the various steps of the pipeline. Steps with program names in brackets constitute the parts of the mGEMS pipeline. Presented values from mSWEEP and mGEMS binner are the actual results of running the pipeline with the described input.

In the pseudoalignment part of the pipeline (Figure 1), we use our own more efficient and accurate implementation of the pseudoalignment algorithm in kallisto [39], called Themisto, to pseudoalign the sequencing reads against the reference sequences. Themisto is based on using colored de Bruijn graphs to represent the reference sequences and disk storage to control the amount of memory required in constructing the pseudoalignment index. These choices lead to Themisto aligning a similar number of reads per hour as kallisto, while being 70 times faster to load in an example pseudoalignment index consisting of 3682 *E. coli* sequences (28 minutes for kallisto and 0.55 minutes for Themisto; Supplementary Methods). Implementation of the method is described in more detail in Supplementary Methods.

The pseudoalignments from Themisto are used as input to the mSWEEP method [40] to estimate the probabilistic read assignments and whole-sample relative sequence abundances. These values provide the necessary input to the mGEMS binner which assigns the sequencing reads to the bins. Finally, we use the Shovill [41] assembly optimizer for the SPAdes assembler [42,43] to assemble the bins. On an example synthetic mixed sample (the *E. coli* sample with the most reads), the full mGEMS pipeline took 112 minutes to run (Themisto 26 min, mSWEEP 4 min, mGEMS binner 16 min, and Shovill 66 min) using two threads on a laptop computer with two processor cores and 16 gigabytes of memory. C++ implementations of both the mGEMS binner and the Themisto pseudoaligner are freely available on GitHub (https://github.com/PROBIC/mGEMS, MIT license, and https://github.com/algbio/themisto, GPLv2 license).

### Overview of the experiments used in benchmarking mGEMS

We assessed the accuracy and effectiveness of mGEMS by considering data from three genomic epidemiological studies [29,37,38] and by generating a benchmarking dataset of *in vitro* mixed samples with measured DNA concentrations. The *in vitro* dataset was generated by first growing three strains of *E. coli* and three strains of *E. faecalis* separately, resulting in six overnight cultures in liquid medium. Next, the amount of DNA extracted from the overnight cultures was measured and six mixtures, each consisting of three strains of either *E. coli* or *E. faecalis*, with known concentrations of DNA for each isolate, were created. This resulted in a benchmark dataset where the relative abundances of the different strains in each mixture are known. We also generated two additional mixtures where the *E. coli* or *E. faecalis* strains were mixed in 1:1:1 proportions from the liquid culture without measuring the amount of DNA, and the DNA extraction was then performed on these already-mixed bacterial samples. To our knowledge, these benchmarking samples constitute the first published dataset where DNA from three strains of the same species has been mixed with known concentrations, providing an important resource for development of methods aimed at untangling strain-level variation.

In the synthetic mixture experiments, we used sequencing reads from previously published genomic epidemiological studies [29,37,38] as the basis for creating synthetic mixture data. The synthetic mixtures were processed with the mGEMS pipeline, and the output was compared against the benchmark of having non-mixed data available by running the same epidemiological analyses on both the mGEMS output and the non-mixed data. The synthetic experiments presented are: 1) mixing reads from three clones of *E. coli* sequence type (ST) 131 sublineages obtained from a study of multidrug-resistant *E. coli* ST131 strains circulating in a long-term care facility in the UK [37], 2) mixing reads from seven *E. faecalis* STs identified in a study of the population structure of hospital-acquired vancomycin-resistant *E. faecalis* lineages in the UK and Ireland [38], and 3) mixing reads from three *S. aureus* ST22 sublineages from a study of the transmission network of methicillin-resistant *S. aureus* (MRSA) among staff and patients at an UK veterinary hospital [29]. We also provide three different approaches to constructing the reference datasets for the pseudoalignment step: 1) a national (UK) collection of *E. coli* ST131 isolates associated with bacteremia [44], 2) a global collection of all available *E. faecalis* genome assemblies from the NCBI as of 2 February 2020, and 3) a local collection of *S. aureus* sequencing data from the staff members at the veterinary hospital at the earliest possible time point in the same study [29]. A detailed description of the generated experiments and the accession numbers of the isolate sequencing and reference data used is presented in the Methods section.

### Evaluation of mGEMS and mSWEEP on the *in vitro* benchmark data

We first evaluated the accuracy of mGEMS and mSWEEP on the six *in vitro* experimental samples, where the true relative abundances of the three strains in each sample are known. In the *E. faecalis* samples each of the three strains originated from a different multilocus sequence type (MLST) and the measurements were accordingly performed on the level of the MLST grouping [45]. In the *E. coli* samples, the strains originated from sublineages within ST131 as defined in a previous study [44], with one strain from sublineage ST131-A and two from sublineage ST131-C2. In order to distinguish between the two strains from ST131-C2, we further split the strains based on their accessory genomes using PopPUNK [46], which provided us with a grouping where all three strains were split into three separate groups (A-14, C2-4, and C2-6), enabling us to differentiate between them with mSWEEP and mGEMS.

We assessed the accuracy of mGEMS by comparing the results of SNP calling from a hybrid long+short-read assembly obtained from a single-colony derived sample of the strains used in the *in vitro* mixed experiments with calling the SNPs from an assembly obtained by processing the mixed experiment samples with the mGEMS pipeline. The results of the SNP calling are highly similar in both datasets (Figure 2 panels a and b) with the exception of the *E. coli* ST131-C2-6 strain from the experiment labelled “Exp 2 *E. coli*”. In this experiment, the sample consisted of equal amounts of DNA from the ST131 C2-4 and C2-6 strains and a small amount of ST131-A-14, causing some confusion between the reads originating from the closely related C2-4 and C2-6 strains which resulted in a difference between the observed and expected SNP counts.

**Figure 2.**
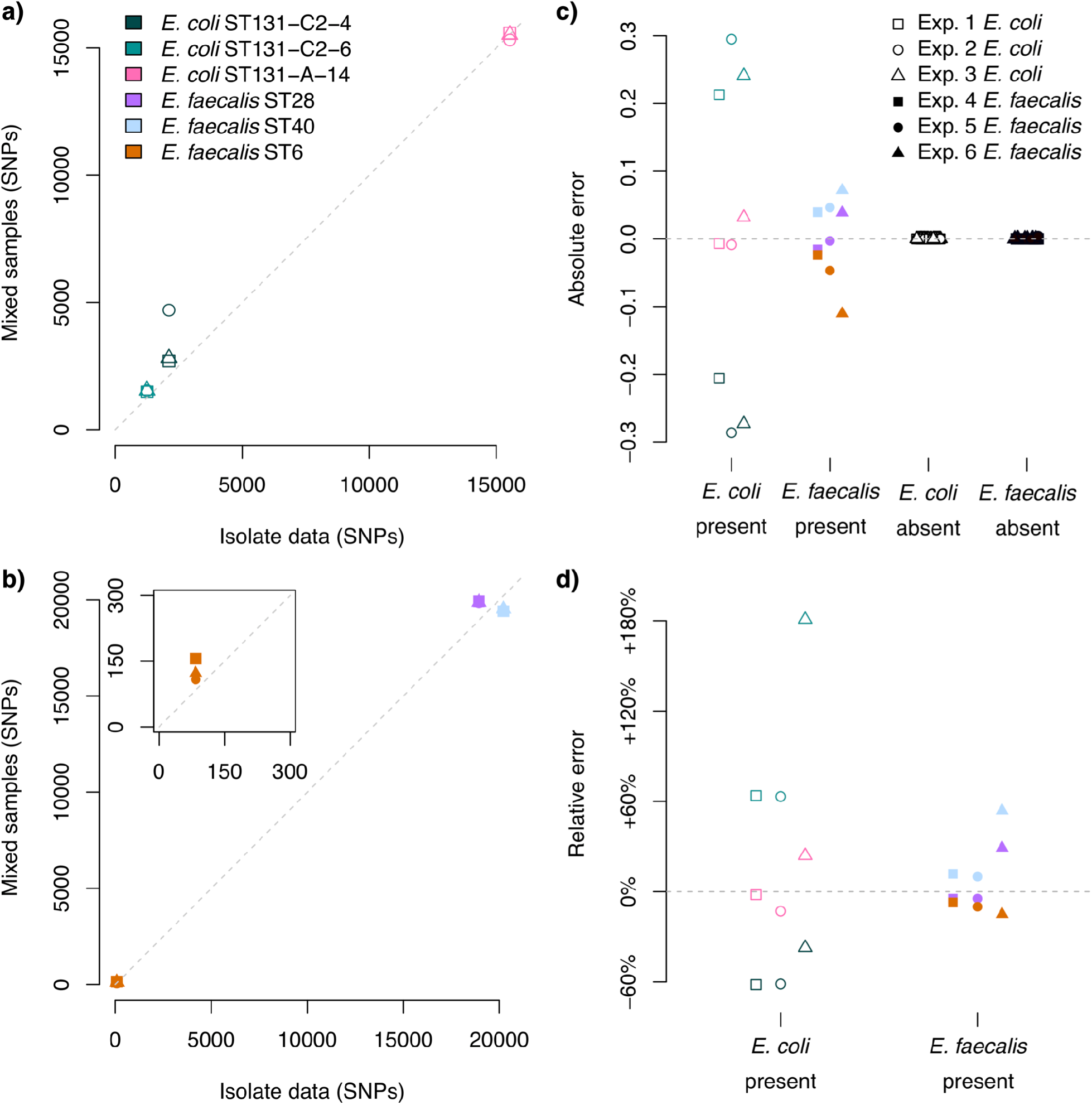
Evaluating mGEMS and mSWEEP on the *in vitro* benchmark data. Panels **a)** *E. coli* and **b)** *E. faecalis* compare the results of SNP calling from the isolate sequencing data (horizontal axis) against the results of SNP calling from the mixed samples with the mGEMS pipeline (vertical axis). The subplot in panel **b)** contains a zoomed-in view of the points around the origin. Panels **c)** and **d)** compare the abundance estimates from mSWEEP to the ground truth relative abundances. Panel **c)** shows the absolute difference between the estimates from mSWEEP and the true abundance. The values shown are split into *E. coli* and *E. faecalis* lineages truly present in the samples, and lineages truly absent. Panel **d)** shows the relative error in the truly present lineages.

Similarly, the mSWEEP relative abundance estimates for both the *E. coli* and *E. faecalis* samples correspond well with the true values when measured by both absolute and relative error (Figure 2 panels c and d, respectively). Slightly higher errors were observed in the estimates for the *E. coli* ST131 C2-4 and C2-6 strains when compared to the estimates for the *E. coli* ST131 A-14 strain and all three *E. faecalis* strains. Akin to the results of SNP calling with mGEMS, these differences in the relative abundance estimates are likely a result of using the highly detailed *E. coli* within-ST clustering, which is significantly harder to differentiate than the between-ST clustering used for *E. faecalis*. Regardless, in all cases, there are no false positive or false negative detections of lineages reported in the mSWEEP relative abundance estimates.

### SNPs from synthetic mixtures match SNPs called from isolate data

In the first synthetic mixture benchmark, we compared the accuracy of SNP calling with the snippy software (v4.4.5) [47] from the bins obtained by processing the abundance estimation results from the mixed samples with the mGEMS binner with the results of the same analyses from the isolate sequencing data (Figure 3). In the *E. coli* and *E. faecalis* experiments (Figure 3 panels a and b, respectively), the SNPs were called from assembled contigs while in the *S. aureus* experiment (Figure 3 panel c), we called the SNPs directly from the sequencing reads because calling the SNPs from the contigs resulted in poorer performance (Supplementary Figure 1). In all experiments, the SNPs called from the mixed samples closely resemble the results of isolate sequencing data in both the samples that are similar and dissimilar to the reference sample. Although in the *E. coli* experiment mGEMS produced slightly more SNPs on average, the results were consistently higher for all samples and did not affect the results of the analyses presented further in this article.

**Figure 3.**
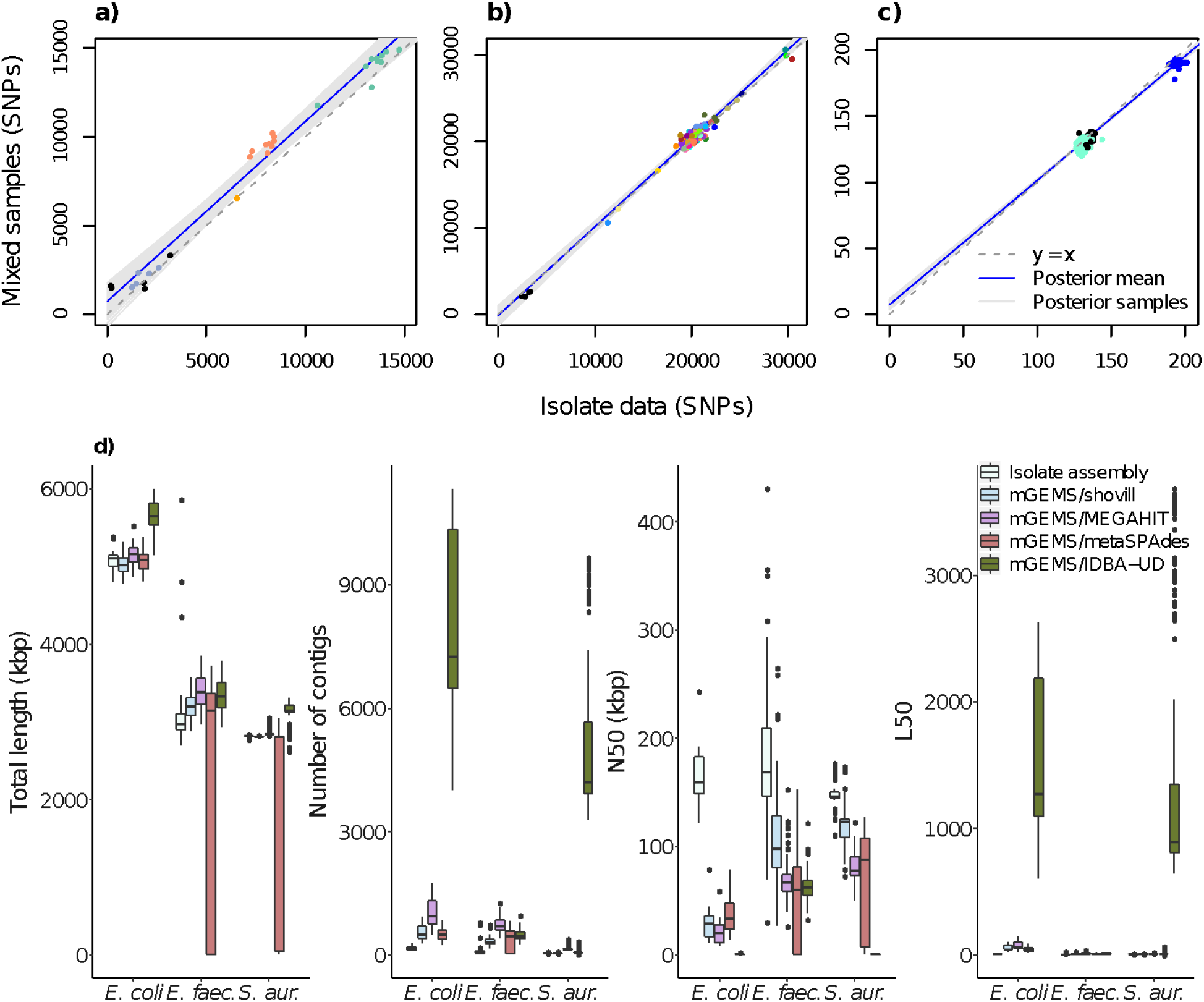
Comparing mGEMS and synthetic mixtures with isolate sequencing data. Panels **a**, **b**, and **c** compare the results of SNP calling from mixed samples with the mGEMS pipeline against the results from isolate sequencing data. Panel **d** compares reference-free assembly statistics from mGEMS pipeline with different assemblers against the results from assembling the isolate sequencing data with Shovill. The results in panel a are for the *E. coli* ST131 isolates, panel **b** the *E. faecalis* isolates, and panel c the *S. aureus* ST22 isolates. In panels **a** and **b**, SNPs were called from contigs after assembling the reads. In panel **c**, the SNPs were called directly from the reads. Points are colored according to the lineage within the species. The dashed gray line represents a hypothetical perfect match between the binned and isolate reads. The blue line is the posterior mean while the shaded area contains the 95% posterior credible region calculated from 10 000 posterior samples from a Bayesian regression model with the SNPs from the binned reads as the response and the SNPs from the isolate sequencing data as the sole explanatory variable. In panel **d**, the boxes are colored according to the type of assembly. The presented statistics are the summed lengths of all contigs (total length), the number of contigs, the sequence length of the shortest contig at 50% genome length (N50), and the smallest number of contigs whose sum of lengths is at least 50% of the genome length (L50).

We suspected that the observed differences in the SNP counts may have been caused by issues in the sequence assembly due to mGEMS allowing a read to belong to multiple bins, which results in variable coverage between the regions with and without the clade-specific SNPs. We tested this assumption by replacing the Shovill assembler in the mGEMS pipeline with metagenomic assemblers, which naturally handle variable coverage. Using the metagenomic assemblers marginally improved the results in some of the experiments (Figure 3 panel d, Supplementary Figure 2). However the improvements were not drastic enough to decisively confirm our suspicions about the accuracy of the SNP calling being limited by the choice of assembler. We did observe that when measured by reference-independent assembly statistics (sum of all contig lengths, total number of contigs, sequence length of the shortest contig at 50% genome length N50, and the smallest number of contigs whose sum is at least 50% of the genome length L50), the statistics obtained from the standard configuration of mGEMS with the Shovill assembler resemble those from isolate sequencing data.

We further assessed the accuracy of the called SNPs by fitting a Bayesian linear regression model to the same SNP data with the isolate results as the sole explanatory variable and the results from the bins or the metagenomic assemblers as the response variable (Figure 3 and Supplementary Figure 2) using the brms R package [48–50]. In both the *E. coli* ST131 sublineage and the *E. faecalis* experiments, the 95% posterior credible interval for the slope from mGEMS with all assembler choices except metaSPAdes contains the correct value of 1.0. The *S. aureus* experiments produce worse 95% credible intervals for the slope compared to the *E. coli* and *E. faecalis* experiments with none of the intervals containing the correct value. However, the regression model is not well suited to analysing the *S. aureus* samples since the number of SNPs between the strains is minimal (0-10 SNP differences within the lineages) and there are only three lineages, which translates poorly to finding a linear relationship.

### Split-*k*-mer comparison between isolate reads and mGEMS bins in synthetic mixtures

We also examined the accuracy of the mGEMS binner without assembling by using the split *k*-mer analysis provided by the SKA software (v1.0, [51]). In a split-*k*-mer analysis, each nucleotide in the read is flanked by two *k*-mers. The nucleotide in the middle position plus the flanking *k*-mers constitute a single split-*k*-mer. If the split*-k*-mers are calculated for all nucleotides in two samples, they can be used to compare the samples on the basis of matching or mismatching split-*k*-mers or to call SNPs by comparing two split-*k*-mers where the flanking *k*-mers match but the nucleotide in between does not.

We first used SKA to call split-*15*-mer-SNPs in the three reference sequences from the binned sequencing reads, and calculated the difference in the count of SNPs called in the reference sequence between the isolate and the binned reads (Supplementary Figure 3). Since the results in Figure 3 for *S. aureus* were obtained without assembly, there is no notable difference when compared to the SKA results.

However, the SKA results for *E. coli* and *E. faecalis* contain fewer SNPs called from the binned reads, implying that binning with mGEMS acts as filtering for the sequencing data, since the results from the assemblies display no stark differences.

In our next assessment, we performed pairwise comparisons within the separate sets of 1) all isolate reads, and 2) the binned reads. First we called the split-*15*-mer SNPs pairwise between all samples containing the isolate reads, and pairwise between all samples containing the binned reads. We then calculated the differences in the pairwise SNP counts obtained from the isolate reads and the binned reads. Secondly, we performed the same pairwise analysis but instead of the split-*15*-mer SNP counts we looked at the numbers of split-*15*-mers that either were the same (matching) or different (mismatching) between each pair of samples. The results from these two comparisons (Supplementary Figure 4) show more discrepancy than the earlier results considering only SNPs called in the reference genome (Figure 3), but the pairwise SNP counts are still relatively well preserved in all three species.

### Phylogenetic analysis of *E. coli* ST131 sublineages in a long-term care facility with synthetic mixtures

We used a set of 30 multidrug-resistant *E. coli* ST131 strains sequenced from the residents of a long-term care facility in the UK [37] to produce a total of 10 synthetic mixed samples. Each sample was the result of mixing isolate sequencing data from three *E. coli* ST131 sublineages (one from each of the main lineages A, B, or C) together. We attempted to preserve the potential sequencing errors and biases by using all available reads from each of the isolate samples.

We applied the mGEMS pipeline to the 10 synthetic mixed samples with a national (from the UK) collection of *E. coli* ST131 strains as the reference data [44], and used RAxML-NG (v0.8.1) [52] to infer a phylogenetic tree from both assemblies obtained from the isolate sequencing data (ground truth) and the assemblies from the mGEMS pipeline. Comparisons of these two trees (Figure 4), show that the overall structure of the trees is highly similar, with the deep branches within the tree well reconstructed and differences in the tree topology appearing only at the very recent short branches.

**Figure 4.**
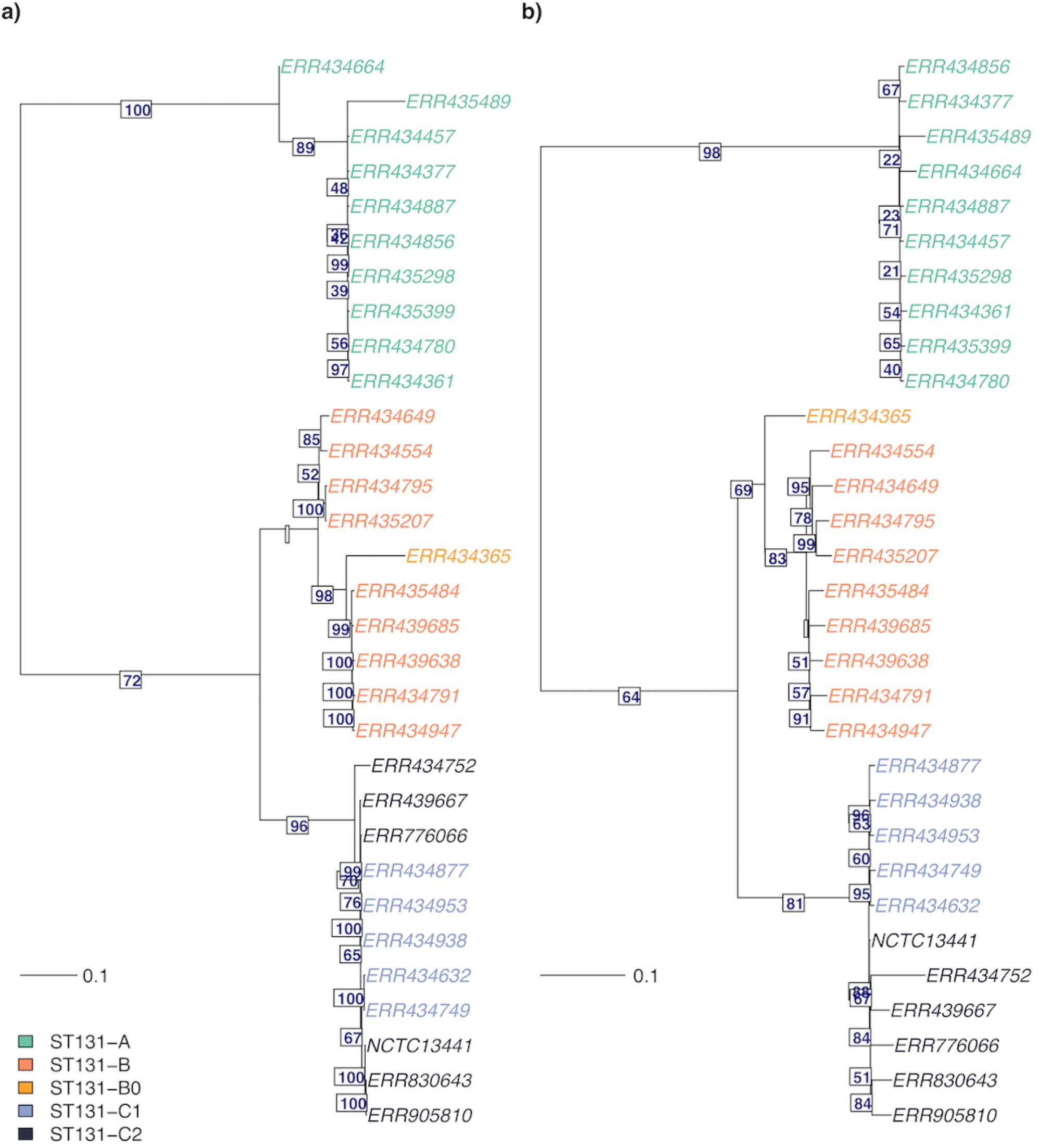
Midpoint-rooted maximum likelihood trees from core SNP alignment of *E. coli* ST131 strains. The phylogeny in panel **a** was constructed from isolate sequencing data from 30 *E. coli* ST131 strains, and the phylogeny in panel **b** with the mGEMS pipeline from 10 synthetic plate sweep samples, each mixing three isolate samples from the three main ST131 lineages (A, B, and C; one strain from each per sample). Both phylogenies were inferred with RAxML-NG. Numbers below the edges are the branch support values from RAxML-NG for the next branch. Leaves are coloured according to the *E. coli* ST131 sublineage (A, B, B0, C1, or C2), and branch lengths in the tree scale with the mean number of nucleotide substitutions per site on the respective branch (GTR+G4 model). Leaves are labelled with the ENA accession number and the leaf labelled *NCTC13411* corresponds to the reference strain used in calling the core SNPs.

### Population structure of nosocomial *E. faecalis* infections in the UK from synthetic mixtures

Our next experiment was performed on sequencing data from bloodstream-infection-associated *E. faecalis* strains with a high prevalence of vancomycin resistance circulating in hospitals in the UK [38]. In this experiment, we mixed isolate sequencing data from seven distinct *E. faecalis* STs [45], producing a total of 12 synthetic mixed samples with seven lineages present in each. Each synthetic mixed sample included all sequencing reads from the mixed isolate sequencing data similarly to the *E. coli* experiment. We used a global collection of *E. faecalis* strains (all *E. faecalis* genome assemblies submitted to the NCBI as of 2 February 2020) as the reference data for the mGEMS pipeline, and again inferred the phylogenies for assemblies from both the isolate sequencing data and the results of the mGEMS pipeline. The more complex structure of these phylogenies was compared by plotting the two phylogenies against each other in a tanglegram (Figure 5). Apart from a few structural mismatches in branches with poor bootstrap support values in both phylogenies (indicating uncertainty in the structure to begin with), the tree structure is strikingly well-recovered from the binned reads.

**Figure 5.**
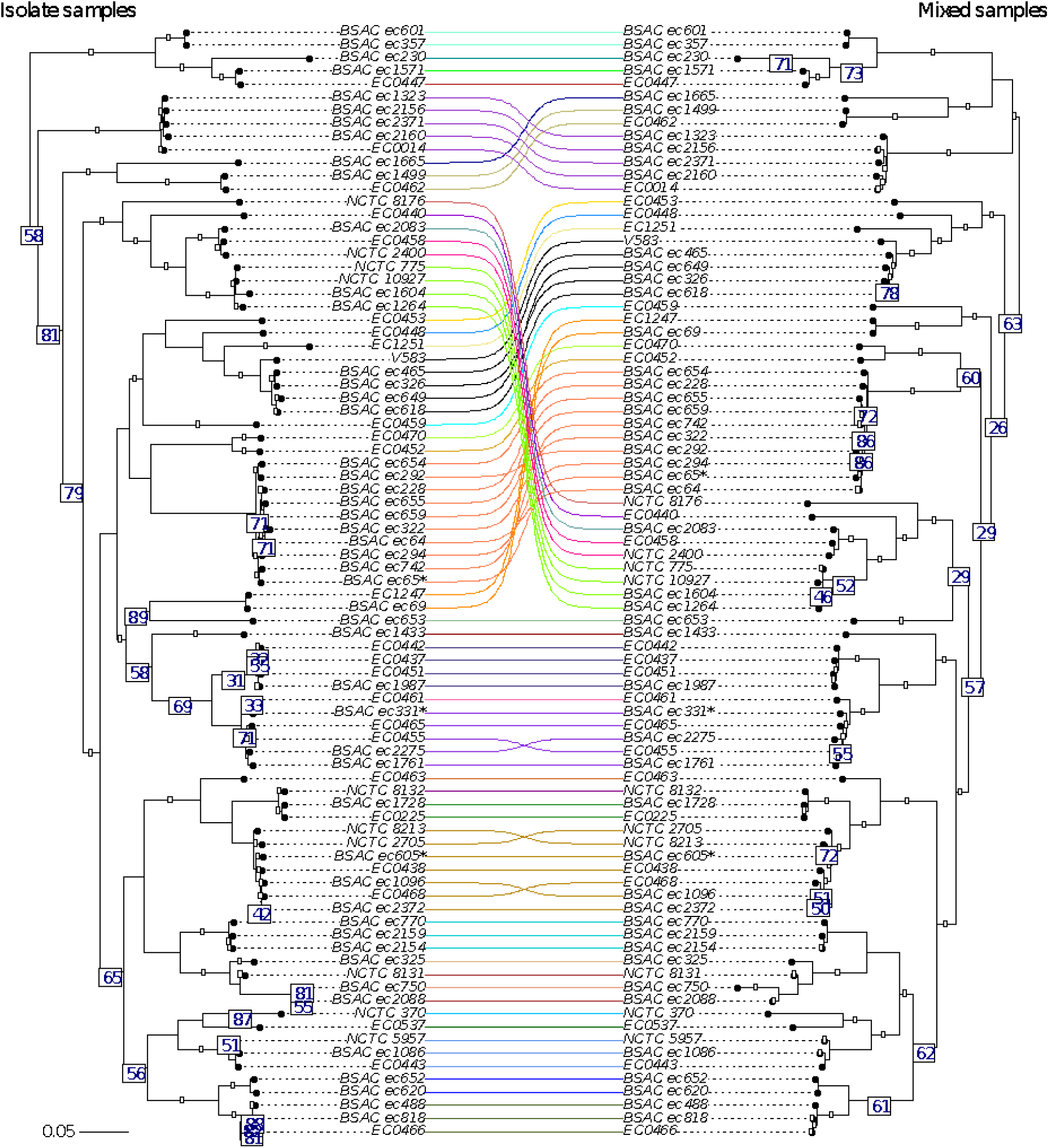
Tanglegram of two midpoint-rooted maximum likelihood trees from core SNP alignment of *E. faecalis* strains. The phylogeny labelled *Isolate samples* (left side of the tree) was inferred with RAXML-NG from assembling the isolate sequencing data from 84 *E. faecalis* strains. The phylogeny labelled *Mixed samples* (right side of the tree) was inferred from 12 synthetic mixed samples, each containing sequencing data from seven different *E. faecalis* STs randomly chosen from the isolate sequencing data. Numbers below the edges indicate bootstrap support values from RAxML-NG for the next branch towards the leaves of the tree. Only support values *less than* 90 are shown. Branches are coloured according to the *E. faecalis* STs, and branch lengths in the tree scale with the mean number of nucleotide substitutions per site on the respective branch (GTR+G4 model). Leaves are labelled with the strain name from NCBI and the leaf labelled *V583* corresponds to the reference strain for calling the core SNPs.

In fact, the tree inferred with the mGEMS pipeline has better bootstrap support values in the lower parts of the tree, suggesting that using mGEMS provides a better phylogeny than using the isolate sequencing data alone. We suspect this improvement in the bootstrap support values was caused by contamination in the isolate sequencing data for BSAC ec750, which produces an assembly 5.8Mb long — nearly twice the length of the reference *E. faecalis* strain V583 (3.2Mb). Similar changes in the bootstrap support values and additional structural changes occur in the parts of the tree containing the isolates BSAC ec294 and BSAC ec655 which both produce abnormally long assemblies (4.8Mb and 4.4Mb, respectively). The assembly lengths for both the isolate and mGEMS-binned sequencing reads are provided in Supplementary Table 1.

### Methicillin-resistant *S. aureus* transmission patterns among staff and patients at a veterinary hospital from synthetic mixtures

In our last experiment, we used a dataset containing three *S. aureus* ST22 sublineages (called clade 1, clade 2, and clade 3) circulating among the staff and patients at a veterinary hospital in the UK [29] and separated by less than 150 SNPs. Because of the minimal differences between the clades, and a lack of isolates from these specific clades in published sources, we decided to use the isolates from the temporally first sample from the staff members as the reference data (representing a local reference collection). We separated the reference isolates from our experiment cases, which consist of all samples sequenced after the reference isolates, and proceeded to mix the remaining isolate sequencing data together. We generated a total of 312 synthetic mixed samples, each containing the sequencing data from three isolate samples from each of the three clades. Because the numbers of samples in each clade were not equal, the data from some of the isolate samples were included in multiple mixed samples. Since we wanted to represent each isolate with only a single instance in the phylogeny, we randomly chose one corresponding bin from mGEMS as the representative for an isolate that was included in multiple mixed samples.

The phylogenies in Figures 5 and 6 were inferred with RAxML-NG (v0.8.1, [52]) from the results of the mGEMS pipeline. We plotted the subtrees of the overall phylogeny separately for the clade 1 isolates (Figure 6) and clade 2 and 3 isolates (Figure 7) without changing the underlying tree structure. Phylogenies inferred from the isolate sequencing data using the same pipeline are available in Supplementary Figure 5 and Supplementary Figure 6. In the original study [29], Staff member A was inferred as having introduced the MRSA strain from Clade 1 into the veterinary hospital. In our phylogeny (Figure 6), Staff member A’s initial samples (timepoints labels 1 and 2) are indeed contained at the root of the tree inferred from the mGEMS pipeline, although the placement of the strains further up the tree vary when compared to the results presented in the original study. The original study performed manual quality control of the SNP data by removing transposable elements which was not replicated in our experiment, possibly explaining some of the observed differences between the tree structures. The phylogenies for clades 2 and 3 (Figure 7) follow the results of the original study more closely with most subclades found in both the isolate and the mixed sample phylogenies. Importantly, in all three clades no assembly from the mGEMS pipeline was assigned to the wrong clade in the phylogeny despite the minimal distances between the clades.

**Figure 6.**
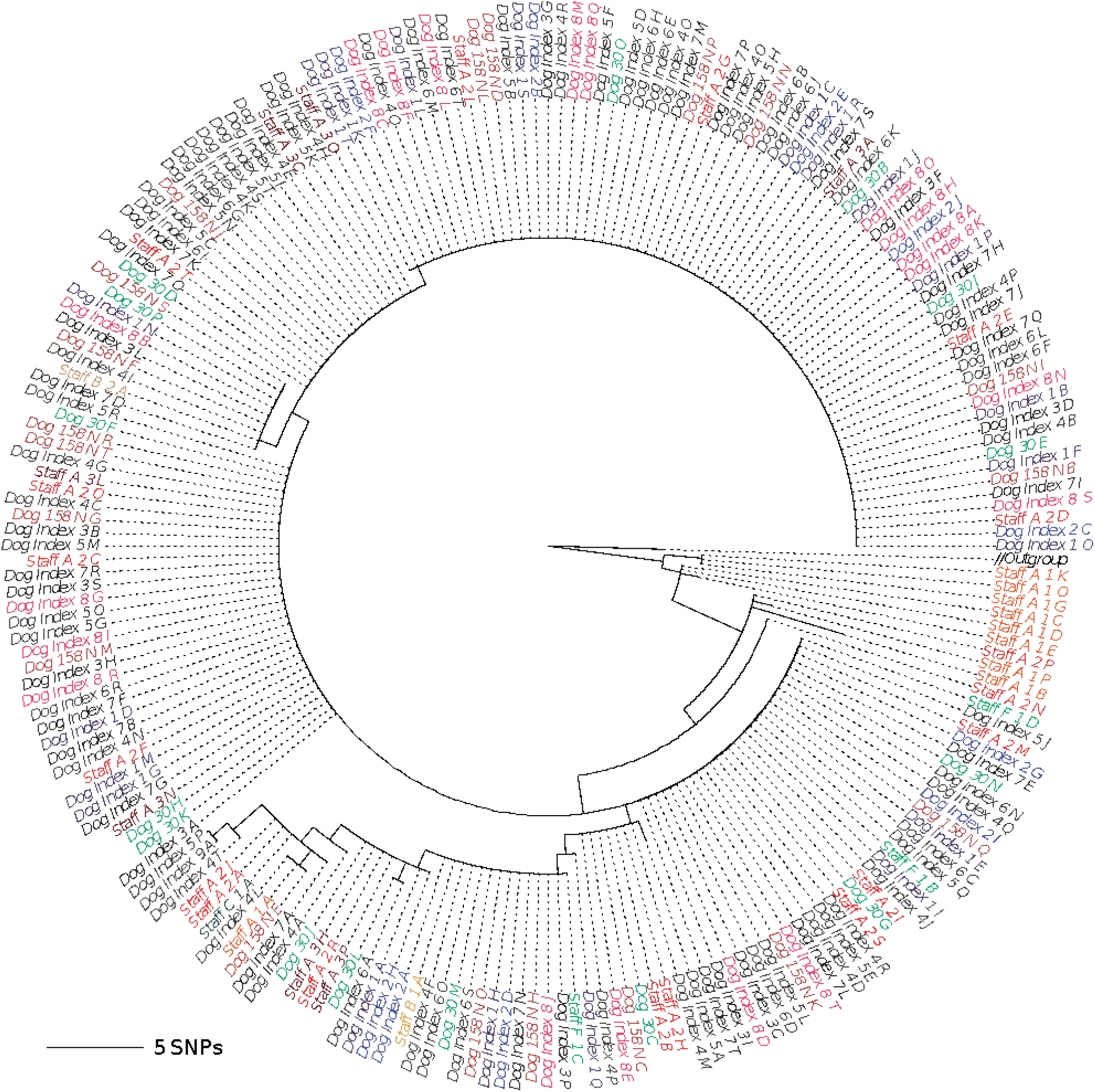
Midpoint-rooted maximum likelihood tree from core SNP alignment of *S. aureus* ST22 showing strains from a single lineage within the sequence type. The phylogeny was inferred from a combined set of assemblies from 60 isolate sequencing samples (leaves labelled Staff A-G 1 A-T, corresponding to the temporally first samples from each staff member) and 312 assemblies obtained from the mGEMS pipeline applied to synthetic mixed samples of sequencing data from each of the three different *S. aureus* ST22 clades (1, 2, and 3). Only strains from clade 1 are displayed in the tree, with the branch labelled *Outgroup* leading to the collapsed clades 2 and 3. The mixed samples were produced from the isolate sequencing data collected from the patients, or from the staff members after the first sampling time. Branch labels are coloured according to the plate the isolate sequencing data was picked from. Branch lengths in the phylogeny scale with the mean number of SNPs obtained by multiplying the mean nucleotide substitutions per site on the respective branch (GTR+G4 model) with the total number of alignment sites. Leaves are labelled with the format: staff or patient, a letter indicating the donor, plate number (ascending in time), and a letter indicating the colony pick id.

**Figure 7.**
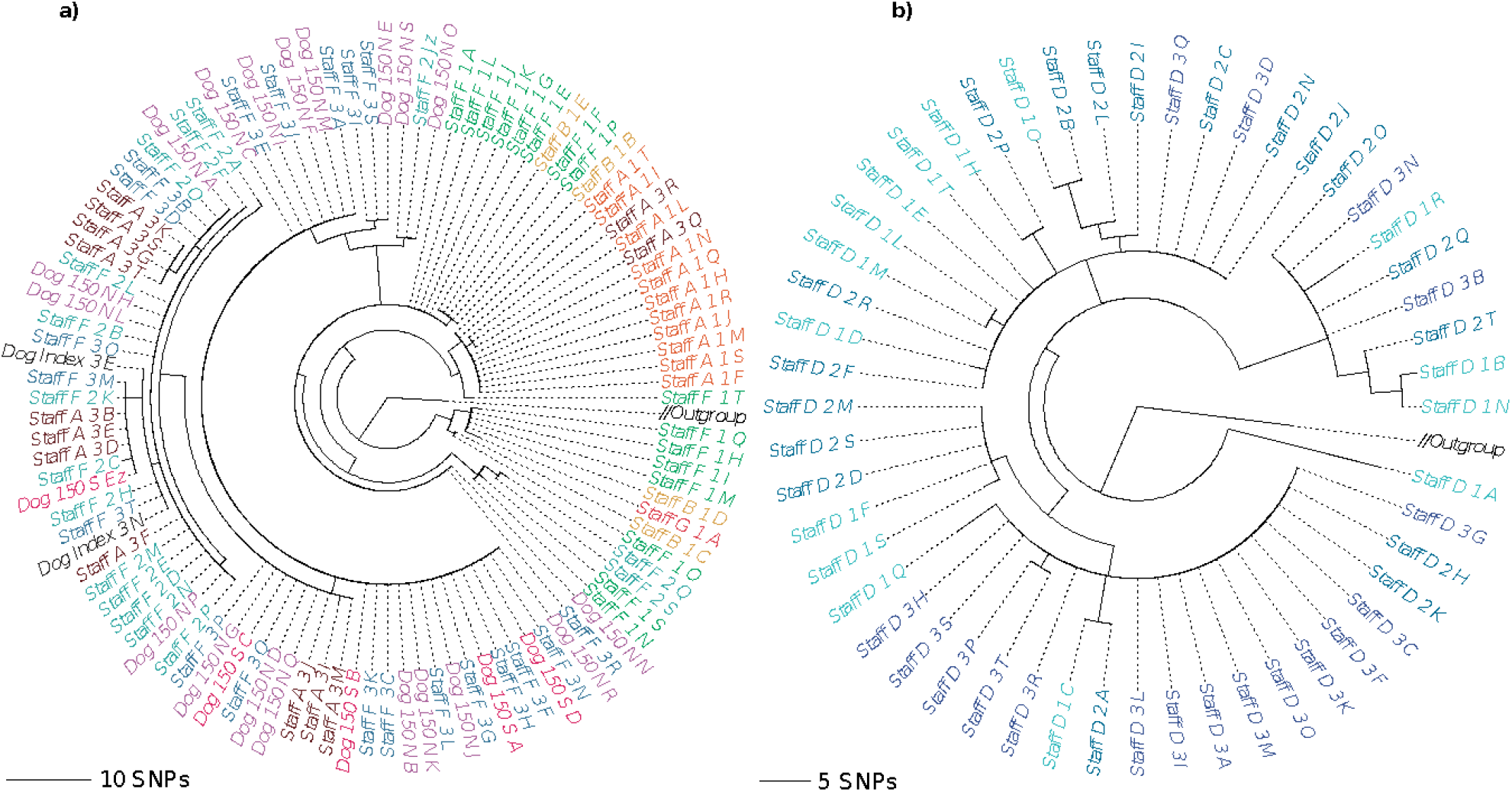
Midpoint-rooted maximum likelihood trees from core SNP alignment of *S. aureus* ST22 showing clade 2 and clade 3 strains. The underlying phylogeny is the same as in Figure 6. The phylogeny in panel **a** contains the clade 2 strains, and panel **b** the clade 3 strains. Branches leading to clade 1 and clade 3 (panel **a**), or clade 1 and clade 2 (panel **b**), labelled *Outgroup* in both panels, were collapsed. Branch labels are coloured according to the plate the isolate sequencing data was originally picked from with darker shades indicating later sampling times. Branch lengths in the phylogeny scale with the mean number of SNPs obtained by multiplying the mean nucleotide substitutions per site on the respective branch (GTR+G4 model) with the total number of alignment sites. Leaves are labelled with the format: staff or patient, a letter indicating the donor, plate number (ascending in time), and a letter indicating the colony pick id.

## Discussion

Adopting a plate-sweep approach, where DNA from the individual bacteria growing on the same plate is prepared and sequenced as a single library, shows clear promise in reducing the amount of manual and costly laboratory work that has been identified as an emerging bottleneck for epidemiological analyses at many public health laboratories [6]. In this article, we have introduced the mGEMS pipeline, which includes novel pseudoalignment and read binning methods, for genomic epidemiological analyses of plate sweeps. Our pipeline provides means to accurately recover the genomes, or corresponding sequencing reads, from mixed samples with extremely closely related strains separated by less than a few dozen SNPs. In these settings, where the differences between the strains are at or under the sequence type level, isolate sequencing is traditionally required to draw epidemiological conclusions.

Using both samples based on synthetically mixed reads, as well as experimentally generated benchmark samples mixing bacterial DNA and strains, we have shown that with mGEMS we can robustly infer the same conclusions from plate sweeps that can be inferred from single-isolate sequencing data. Additionally, since mGEMS relies on modelling counts of pseudoalignments against grouped reference sequences, the inclusion of the alignment step causes the pipeline to also acts as quality control for sequencing reads from samples that inadvertently contain multiple lineages or contamination, which can disrupt downstream analyses like SNP calling [53]. In analysing sequencing data from closely related mixed samples our pipeline reaches accuracy levels likely constrained by technical variation in the sequencing data and limitations in assembling sequencing data with variable coverage. To our knowledge, mGEMS is the first tool capable of reliable recovery of the full strain variety in complex mixed samples.

mGEMS demonstrates the power of plate sweep sequencing in genomic epidemiology and enables a change in the currently dominant framework that confers multiple benefits over both whole-genome shotgun metagenomics and isolate sequencing. Studies of the population structures of opportunistic pathogens have revealed extensive strain-level within-host variation [20,22,26,54,55] with adverse implications for transmission analyses relying solely on isolate sequencing [30,56] and colony pick based longitudinal studies reporting the absence or re-emergence of strains in a host [29,37,57] or antimicrobial profiles [27,58]. While whole-genome shotgun metagenomics solves these issues to some extent [34,59], the culture-free nature suffers from issues with both bacterial and host DNA contamination particularly affecting the sensitivity for detecting strains in low abundance [28,32,33,60,61]. Using mGEMS in conjunction with plate sweep sequencing data avoids these issues altogether, paving way for more representative studies of pathogen population structure and providing higher-resolution data for more complex models of transmission dynamics incorporating within-host variation and evolution [62–64].

Since our method relies on available single-clone genomic reference data and plate cultures of the bacteria to sequence them at a sufficient depth for assembly, it obviously cannot be applied to the study of uncharacterized or unculturable species. However, culture media do exist for most human pathogens of public health relevance [36] or can be developed for some of the until recently unculturable bacteria [65–67]. Moreover, the availability of single-clone bacterial genome sequences is still increasing at a high rate, such that for many species or lineages plenty of sufficiently representative reference sequences would be available [68,69]. In these cases, the drastic reduction in the costs of library preparation, and the better capture of the underlying genomic variation between closely related bacteria in a set of mixed samples provided by mGEMS is extremely valuable. We hope that by enabling significant streamlining of the process of producing data for public health genomic epidemiology, our approach inspires both applications and further method development within this exciting research area.

## Conclusions

We have developed the mGEMS pipeline for performing genomic epidemiological analyses from mixed samples containing multiple closely related bacterial strains. The two crucial novel enabling aspects introduced in this paper are the mGEMS read binner and the Themisto pseudoaligner. The mGEMS binner is a binning method based on turning probabilistic assignment of sequencing reads to reference lineages, while the Themisto pseudoaligner is a high-throughput exact pseudoaligner for short-read sequencing data that features external memory construction for compressed coloured de Bruijn graphs for scalability, providing significant runtime savings over conventional pseudoalignment. mGEMS addresses several major issues related to the cost, applicability, and sensitivity of the current approach in genomic epidemiology and enables entirely new types of analyses using mixed samples without sacrificing accuracy.

## Methods

### mGEMS workflow

Our pipeline for performing genomic epidemiology with short-read sequencing data from mixed samples, mGEMS, requires as input the sequencing reads and a reference database representing the clonal variation in the organisms likely contained in these reads. The reference database must additionally be grouped accordingly into clonal groups representing lineages within the species. We used either the multilocus sequence types (*E. faecalis* experiments) or sublineages within the sequence types (*E. coli* and *S. aureus* experiments) as the clonal grouping. With these pre-processing steps performed, the first step in the mGEMS pipeline is to pseudoalign [39] the sequencing reads against the reference database using our scalable implementation of (exact) pseudoalignment with the Themisto software (in this article we used v0.1.1 with the optional setting to also align the reverse complement of the reads enabled). The pseudoalignments and the clonal grouping are then supplied as input to the mSWEEP software (v1.3.2; doi: 10.5281/zenodo.3631062, [40], with default settings) which estimates the relative sequence abundances of the clonal groups in the mixed sample. Consequently, mSWEEP produces a probabilistic assignment of the sequencing reads to the different reference clonal groups. This tentative assignment is subsequently processed by the mGEMS binner (v0.1.1, default settings), which assigns the sequencing reads to bins that correspond to a single reference clonal group — with a possibility for a sequencing read to belong to multiple bins. As the last step, the bins are (optionally) assembled with the shovill (v0.9.0, with default settings, [41]) assembly pipeline. mGEMS and Themisto are freely available on GitHub (https://github.com/PROBIC/mGEMS and https://github.com/algbio/themisto).

### Reference data for mSWEEP and mGEMS

We used assemblies of three different sets of sequencing data as the reference for the three different experiments presented. The three different reference datasets represent a local (*S. aureus* experiment, [29]), a national (*E. coli*, [44]), and a global collection (*E. faecalis* downloaded from the NCBI) of strains from these species. Accession numbers and multilocus sequence types for the reference data are available in Supplementary Table 2 accompanied with rudimentary assembly statistics from both the isolate sequencing data and the assemblies from the mGEMS pipeline. In each experiment, we only aligned against the reference sequences from the relevant species.

In the *E. coli* experiments, our collection of 218 *E. coli* ST131 isolates originated from the British Society for Antimicrobial Chemotherapy’s bacteraemia resistance surveillance program and were originally isolated from 11 hospitals across England [44]. These isolates were assigned to six ST131 sublineages (A, B0, B, C0, C1, or C2) as described in a previous study [44]. As the reference sequence for calling the SNPs in building the phylogeny, we used the ST131 strain NCTC13441 (European Nucleotide Archive [ENA] sequence set UFZF01000000).

For the *E. coli* experiment where the six sublineages were further split, the split was generated by clustering the sequences with PopPUNK (v2.3.0, [46]) with the BGM model using four components and then performing the subsequent refinement step. The resulting PopPUNK clustering was combined with the sublineage numbering by concatenating the two together. This clustering is included in Supplementary Table 2.

The global collection of *E. faecalis* reference data was obtained by downloading all available *E. faecalis* assemblies (1484 as of 2 February 2020) from the NCBI, which were assigned to STs with the mlst software (v2.18.1) [45,70,71]. Sequence type could not be determined for 177 assemblies. These were discarded, leaving a total of 1307 assemblies assigned to 203 distinct sequence types. We used the ST6 strain V583 [72] as the reference for SNP calling (NCBI RefSeq sequences NC_004668.1-NC_004671.1).

The *S. aureus* reference data were obtained from the same study as the experiment data [29]. We used shovill (v0.9.0, [41], with default settings) to assemble the isolate sequencing reads from the first sampling of the staff members at the veterinary hospital, and assigned the assembled sequences to the ST22 sublineages according to the information provided in original study [29]. The reference sequence used in calling the SNPs was the ST22 strain HO 5096 0412 ([73], ENA sequence HE681097.1)

If the reference sequence in any of the experiments consisted of multiple contigs, we concatenated the contigs together by adding a 100-base gap between them. The final reference file that was used as input for Themisto indexing was produced by concatenating all reference sequences processed in this way together.

### *In vitro* mixture experiment setup

We first generated reference genomes for the three *E. coli* and three *E. faecalis* strains used in the *in vitro* benchmarking of mGEMS. In order to obtain as accurate reference genomes as possible, we combined short-read Illumina sequencing data (Supplementary Table 3) with long-read Oxford Nanopore sequencing data.

In the first mixture experiment, single colonies of each strain were grown up overnight in liquid medium (LB broth [Sigma-Aldrich] for *E. coli* and brain heart infusion broth [Fluka Analytical] for *E. faecalis*), and DNA extracted for short-read sequencing. The DNA concentration was quantified using the Qubit system (Invitrogen) and purified DNA, diluted to 30 ng/μL, from the three strains of *E. coli* and the three strains of *E. faecalis* were used to prepare three different mixtures with varying ratios (1:1:1, 1.4:1.4:0.2, and 2.2:0.4:0.4; Table 1). These mixtures were then prepared for Illumina sequencing, and analysed as described earlier in the manuscript. All sequencing data generated for these mixture experiments is available under BioProject PRJNA720284.

**Table 1.** Mixture concentrations in the six *in vitro* experiment samples. The column labels contain the sample identifiers and lineages for the strains that were mixed in each of the six experiments, represented by the rows of the table.

|  |  |  |  |
| --- | --- | --- | --- |
|  | **NORM7910041**<br>*E. coli*<br>**ST131-C2-4** | **NORM7911464**<br>*E. coli*<br>**ST131-C2-6** | **NORM7908673**<br>*E. coli*<br>**ST131-A-14** |
| Exp. 1 *E. coli*<br>(1:1:1) | 10 µL | 10 µL | 10 µL |
| Exp. 2 *E. coli*<br>(1.4:1.4:0.2) | 14 µL | 14 µL | 2 µL |
| Exp. 3 *E. coli*<br>(2.2:0.4:0.4) | 22 µL | 4 µL | 4 µL |
|  | **51271926**<br>*E. faecalis* **ST6** | **51271052**<br>*E. faecalis* **ST40** | **51271223**<br>*E. faecalis* **ST28** |
| Exp. 4 *E. faecalis*<br>(1:1:1) | 10 µL | 10 µL | 10 µL |
| Exp. 5 *E. faecalis*<br>(1.4:1.4:0.2) | 14 µL | 14 µL | 2 µL |
| Exp. 6 *E. faecalis*<br>(2.2:0.4:0.4) | 22 µL | 4 µL | 4 µL |

For the second experimental setup, the six reference strains were again grown in single-clone cultures overnight as described above, and 1:1:1 mixtures of the liquid cultures from the three strains per species were made, centrifuged, and used as sample for DNA extraction for short-read sequencing. Unlike in the first experimental setup, in this setup the concentrations of the different strains were not measured with Qubit and thus are not available beyond the initial 1:1:1 mixture of the liquid cultures.

### DNA extraction

DNA was extracted using the MagAttract HMW kit (Qiagen) for Oxford Nanopore sequencing, and the DNeasy Blood and Tissue kit (Qiagen) for short-read sequencing.

### Sequencing

DNA libraries for long-read sequencing were prepared using the Ligation Sequencing Kit SQK-LSK109 (Oxford Nanopore) in combination with the native barcoding kits EXP-NBD104 and EXP-NBD114 (both Oxford Nanopore) according to the manufacturer’s instruction. DNA was sequenced using an Oxford Nanopore GridION on a R9.4.1 flow cell with an input of 337.5 ng. For short-read sequencing, DNA libraries were prepared using the Nextera XT DNA library kit (Illumina) and sequenced on an Illumina NextSeq550 using a mid-output flow cell, 300 cycles and 2×150 bp paired-end set-up.

### Reference genome assembly for the *in vitro* mixed experiments

We performed hybrid assembly from the long and short reads by first assembling only the long-reads and then using the short-reads for error correction. The initial long-read only assembly was created from the raw long-read sequencing data with Flye (v2.8.2, [74]) and polished with quality controlled long reads (QC’d with filtlong v0.2.0; https://github.com/rrwick/Filtlong) using medaka (v1.2.1, https://github.com/nanoporetech/medaka/). The short reads were used for error correction by first quality controlling them with fastp (v0.21.0, [75]) and then using pilon (v1.23, [76]) to perform the error correction on the long-read assembly. This procedure resulted in closed chromosome and plasmid sequences for all six reference strains. The short reads, long reads, and the produced genomes have been submitted and made available in standard repositories (Supplementary Table 3).

### Synthetic mixture generation

We produced our three synthetic mixture sets by synthetically mixing together the isolate sequencing data from distinct lineages in each of the three studies [29,37,38]. In the *E. coli* experiments, we produced 10 mixed samples with one strain from each of the three main ST131 lineages (A, B, or C) in each sample. In the *E. faecalis* experiments, we mixed together seven strains from seven different sequence types to produce a total of 12 mixed samples. The strains included in each sample were chosen at random without replacement in the *E. coli* and *E. faecalis* experiments. The *S. aureus* mixed samples were produced by randomly mixing together one strain from each of the three sublineages with replacement while ensuring that each strain appears at least once. The sequencing data that were used in the reference dataset were not included in any of the experiments. In all three experiment sets, we used all available sequencing data in the mixed samples, resulting in 8-15 million reads in the experiments. Supplementary Table 1 contains the accession numbers and lineage assignments of the isolate sequencing data in each sample, as well as the assembly statistics from both isolate sequencing and the synthetic mixed samples processed with mGEMS.

### Pseudoalignment

We used Themisto (v0.1.1) with the default settings. Themisto is a *k*-mer-based pseudoalignment tool which encodes sets of *k*-mers as a succinct coloured de Bruijn graph. A read is considered to pseudoalign against a reference sequence if at least one *k*-mer of the read is found in the reference, and each *k*-mer of the read is either found in the reference or not found at all in the database of all references. This can be seen as an exact version of the pseudoalignment algorithm implemented by the tool kallisto [39].

The index was constructed using *31*-mers. Themisto does not distinguish between paired-end reads and single reads, so we decided to consider a paired-end read as pseudoaligned only when both fragments pseudoaligned. We have included this functionality for supporting paired-end reads in both the mSWEEP and mGEMS software implementations.

### Abundance estimation and probabilistic read assignment

We used the mSWEEP [40] software (v1.3.2; doi: 10.5281/zenodo.3631062) with default settings. The program was altered to support pseudoalignments from Themisto, and to output the read-level probabilistic assignments to the reference lineages. We also improved the scalability of mSWEEP by parallelizing the abundance estimation part and reducing memory consumption. These alterations have been included in versions v1.3.2 (Themisto and mGEMS support) and v1.4.0 (parallelization and memory usage improvements) of the software.

### Read binning

In order to collect all reads in a mixed sample that likely originate from the same target lineage, we consider a binning strategy that allows associating the same read with multiple reference lineages. We assume that each reference lineage is represented by, at most, only one target sequence in the mixed sample, and that the sets of reference sequences capture the variation in the reference lineages sufficiently to use them as a substitute for the target sequence which may not be included in the reference sequences. In our formal treatment of the task of binning a set of sequencing reads, we define the task in terms of finding *K* subsets (bins), one for each reference lineage *k* = 1,…, *K*, of the full sets of reads *R* = {*r*_1_,…, *r*_*N*_} denoted by *G*_*k*_ ⊂*R* that contain reads likely originating from the target sequence belonging to the reference lineage *k*. The reads assigned to each subset *G*_*k*_ are determined based on read-level probabilities γ_*n,k*_′ 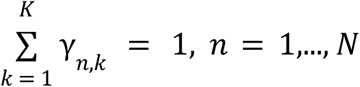 to assign the read *r*_*n*_ into the reference lineage *k* by defining the subsets *G*_*k*_ such that

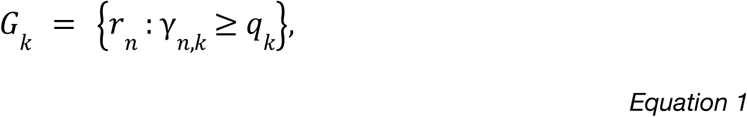

holds for some threshold *q*_*k*_ ∈ [0, 1] which may vary between the lineages *k*. The formulation in Equation (1) has the benefit of allowing the read *r*_*n*_ to possibly belong to several subsets *G*_*k*_, which is an important property for dealing with multiple closely related lineages in the same mixed sample.

In order to find a suitable value for the threshold *q*_*k*_, and to determine the corresponding assignment rule, we consider two binary events: 1) *I*_*n,k*_: the reference lineage *k* generated the read *r*_*n*_, and 2) *j*_*n,k*_: the true nucleotide sequence represented by the read *r*_*n*_ is part of the target sequence belonging to the reference lineage *k*. Knowing the probability of the event *j*_*n,k*_ would directly enable us to assess the plausibility of assigning the read *r*_*n*_ to the reference lineage *k* but its value is difficult to estimate directly. However, we can determine and write down the values of the conditional probabilities *P*[*I*_*n,k*_ = 1 | *j*_*n,k*_ = 0] and *P*[*I*_*n,k*_ = 1 | *j*_*n,k*_ = 1] as

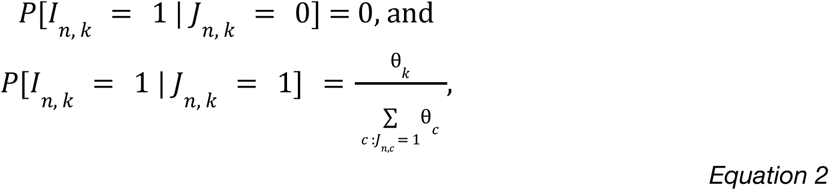

where θ_*k*_ is the proportion of reads from the reference lineage *k*, and 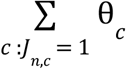 is the proportion of reads from any reference lineages {*c*: *j*_*n,c*_ = 1} which contain the sequence represented by the read *r_n_*. The conditional probabilities in Equation (2) allow us to write the unconditional probability *P*[*I*_*n,k*_ = 1] as

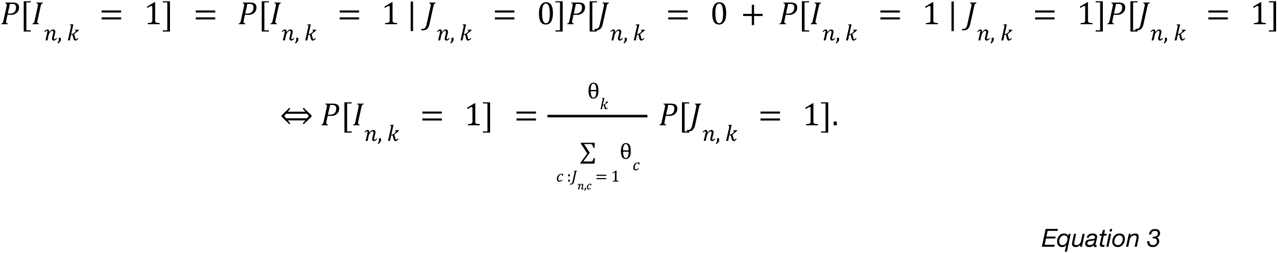

Using the formulation in Equation (3) and the fact that we can approximate 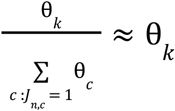 if we assume that the mixed sample is mostly composed of closely related organisms (the denominator 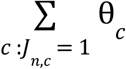 approaches 1), we can rewrite Equation (3) as

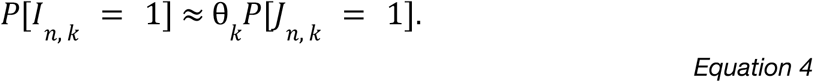

Equations (4) and (3) together imply that if the value of the probability *P*[*I*_*n,k*_ = 1] that the read *r*_*n*_ was generated from the lineage *k* exceeds the relative abundance θ_*k*_ of that lineage in whole sample (*P*[*I*_*n,k*_ = 1] ≥ θ_*k*_), then the value of the probability *P*[*j*_*n,k*_ = 1] that the nucleotide sequence represented by the read *r*_*n*_ is contained in the target sequence from the reference lineage *k* must be “large” (*P*[*J*_*n,k*_ = 1] → 1). This statement about the magnitude of *P*[*J*_*n,k*_ = 1] derives from our assumption that the denominator in Equation (3) is close to 1.

Since we have an estimate of the probabilities *P*[*I*_*n,k*_ = 1] available in the form of the read-level probabilistic assignments γ_*n,k*_ ≈ *P*[*I*_*n,k*_ = 1], we can plug these values in Equation (4) and use the result to derive the assignment rule

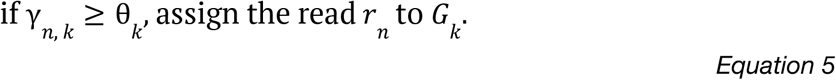

The assignment rule in Equation (5) gives us a way to assess the validity of the statement contained in the probability *P*[*J*_*n,k*_ = 1] which we could not estimate directly.

Because of computational accuracy, we cannot obtain meaningful relative abundance estimates θ_*k*_ for lineages with a relative abundance less than 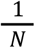 (less than one read from the lineage *k* in the sample). Since there are *K* lineages in total, in the worst-case scenario 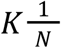 units of the relative abundance fall into this meaningless range. Therefore, only a fraction of the total relative abundance of 1 can be considered to be accurately determined when using computed values of θ_*k*_, and this fraction *d* is determined in the worst-case scenario through the formula

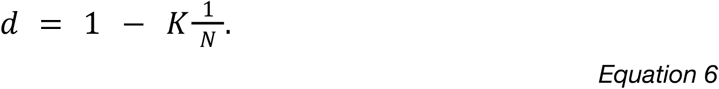

Equation (6) means that when evaluating the validity of the assignment rule presented in Equation (5) with computed values, we have to replace θ_*k*_ with the value *d*θ_*k*_ which depends on the value of *d* in Equation (6). Merging the result from Equations (5) and (6) leads us to the final assignment rule (Equation 7) of

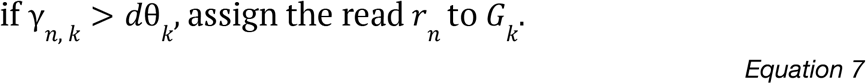

In practice, reads which pseudoalign to exactly the same reference sequences have identical values γ_*n,k*_. The reads can thus be assigned to equivalence classes defined by their pseudoalignments, which enables a speedup in the implementation of the binning algorithm by considering each equivalence class as a single read. Due to this speedup and the computational simplicity of evaluating the assignment rule in Equation (7), the memory footprint of the mGEMS binner is determined by the number of equivalence classes and reference lineages in the input pseudoalignment and the runtime limited by disk I/O performance.

### Genome assembly from mGEMS bins

After binning the sequencing reads in our experiments with the aforementioned assignment rule, we assembled the sequencing reads assigned to the bins using the shovill (v0.9.0, [41], with default settings) assembly optimizer for the SPAdes assembler [42,43]. This step concludes what we in this article call the mGEMS pipeline.

### SNP calling and phylogeny reconstruction

We used snippy (v4.4.5, [47]) to produce a core SNP multiple-sequence alignment from the assembled contigs. Since the *E. coli* and *S. aureus* strains used were from the same sequence type, the core alignment for these two species contained almost the whole genome. After running snippy, we used snp-sites (v2.5.1, [77]) to remove sites with ambiguous bases or gaps from the alignment (*E. coli* experiments only) and then ran RAxML-NG (v0.8.1, [52]) to infer the maximum-likelihood phylogeny under the GTR+G4 model. Since some of the *S. aureus* strains from the same clade were identical, we changed the default value of the minimum branch length parameter in RAxML-NG to 10^−10^ in the *S. aureus* experiments and printed the branch length with eight decimal precision to identify branches of length zero. In all experiments, we ran RAxML-NG with 100 random and 100 maximum parsimony starting trees, and performed 1000 bootstrapping iterations to infer bootstrap support values for the branches. We used the phytools R package (v0.6-99, [78]) to perform midpoint rooting for the tree, and the ape R package (v5.3, [79]) to create the visualizations.

## Declarations

### Ethics approval and consent to participate

This study used only data from published sources, which have sought the appropriate ethical permissions.

### Consent for publication

Not applicable.

### Availability of data and materials

Source code and precompiled binaries (generic Linux and macOS) for both mGEMS and Themisto are freely available in GitHub at https://github.com/PROBIC/mGEMS (MIT license) and at https://github.com/algbio/themisto (GPLv2.0 license). A tutorial describing how to reproduce the synthetic mixed samples, bin the mixed reads, and infer the phylogenies is available in the mGEMS GitHub repository. Accession numbers and other information about the synthetic mixed samples is available in Supplemental Table 1. The reference data used is available from Zenodo (*E. coli* doi: 10.5281/zenodo.3724111, *E. faecalis* doi: 10.5281/zenodo.3724101, *S. aureus* doi: 10.5281/zenodo.3724135), and the accession numbers are available in Supplementary Table 2. Accession numbers for the data used in the *in vitro* mixture experiments is available in Supplementary Table 3.

### Competing interests

The authors declare that they have no competing interests.

### Funding

TM and AH were supported by the Academy of Finland grant no. 310261 as well as the Flagship programme (Finnish Center for Artificial Intelligence FCAI; to JC and AH). TK and JC were supported by the JPI-AMR consortium SpARK (MR/R00241X/1). JC was funded by the ERC grant no. 742158. TK was funded by the Norwegian Research Council JPIAMR grant no. 144501. JA and VM were supported by the Academy of Finland grant no. 309048.

### Authors’ contributions

TM, TK, JC and AH conceived the study, developed the full mGEMS pipeline, and designed the synthetic mixture benchmarking experiments. EH, KH, ØS, TM, JC and AH designed the *in vitro* mixture study. KH, ØS and EH performed the experiments for the *in vitro* study. TM and AH developed the mGEMS binning algorithm. JA and VM developed the Themisto pseudoaligner. TM implemented the mGEMS binner. JA implemented the Themisto pseudoaligner. TM ran the experiments and created the visualizations. TM, TK, JC and AH interpreted the results and wrote the main article. JA wrote the supplementary file describing Themisto. KH, ØS and EH wrote the sections describing the *in vitro* mixture sample generation. All authors participated in reviewing and editing the article and discussed the results.

## Acknowledgements

The authors wish to acknowledge the Finnish Grid and Cloud Infrastructure (persistent identifier urn:nbn:fi:research-infras-2016072533), and CSC – IT Center for Science, Finland for providing computational resources. Silje Lauksund for experimental assistance. Illumina sequencing for the *in vitro* mixture set-up and the Oxford Nanopore sequencing was performed at the Genomics Support Centre Tromsø, UiT The Arctic University of Norway.

## Notes

https://github.com/PROBIC/mGEMS

https://github.com/algbio/themisto

https://zenodo.org/record/3724112

https://zenodo.org/record/3724101

https://zenodo.org/record/3724135

https://zenodo.org/record/4738948

https://zenodo.org/record/4738983

### Competing Interest Statement

The authors have declared no competing interest.

### Summary of Updates

Major revision of the manuscript contents including the addition of a wet-lab benchmark for the method (described under the subheadings "Overview of the experiments used in benchmarking mGEMS" and "Evaluation of mGEMS and mSWEEP on the in vitro benchmark data").

