## Supplementary Methods for "Genomic Epidemiology with Mixed Samples"

### Pseudoalignment in the mGEMS pipeline

In this supplement we describe the pseudoalignment algorithm and implementation used in the mGEMS pipeline. The implementation is called Themisto, and is freely available at https://github.com/algbio/themisto under the GPLv2.0 license. Pseudoalignment is an approximate form of alignment that reports only whether a read matches to a reference sequence or not, without necessarily returning the genomic coordinates of the match. Pseudoalignment can be much cheaper computationally than regular alignment.

#### 1 The pseudoalignment criterion

Our pseudoalignment is based on the pseudoalignment algorithm used in the transcript abundance quantification tool Kallisto [1]. The pseudoalignment criterion we use is defined as follows. Suppose we want to pseudoalign a read against a set of reference sequences $T_1, \ldots, T_m$. The read is considered to pseudoalign against reference $T_i$ if at least one $k$-mer of the read is found in $T_i$ and for each $k$-mer $x$ of the read, one of the following holds:

1. $x$ is a $k$-mer of $T_i$
2. $x$ is not a $k$-mer of any of $T_1, \ldots, T_m$

This criterion closely replicates the pseudoalignment of Kallisto, with the difference that Kallisto uses a heuristic based on the topology of the de Bruijn graph of $T_1, \ldots, T_m$ to skip over some $k$-mers of the read for efficiency. More specifically, if the current $k$-mer is in a non-branching path of the graph, Kallisto skips a number of $k$-mers of the read equal to the distance to reach the next branching node. If the $k$-mers before and after the skip are found in the same reference sequences, the skip is considered valid, and otherwise Kallisto falls back to checking all $k$-mers of the read individually. However, even if the skip is considered valid, it could be the case that a skipped $k$-mer would have affected the result of the pseudoalignment. On the other hand, we implement the described pseudoalignment criterion exactly, and observe a very slight improvement in accuracy compared to using Kallisto's pseudoalignments. The difference in accuracy could also be due to small implementation differences, since we designed our tool around the high-level description in Kallisto's manuscript [1] rather than the source code itself.

#### 2 Implementation overview

The pseudoalignment criterion we have chosen effectively reduces each reference sequence and each read into unorderered sets of $k$-mers. This loses some information, but in turn it allows for more efficient data structures and algorithms. The pseudoalignment could in principle be implemented on top of any data structure for indexing $k$-mer sets.

Indexing $k$-mer sets efficiently is currently a very active field of research [2]. In $k$-mer data structures, each reference sequence is usually given a unique identifier, called the *color* of the sequence. Each $k$-mer is associated with a *color set*, which is defined to be the set of colors of the reference sequences that contain that $k$-mer. The basic query on a $k$-mer data structure is to retrieve the color set of a given $k$-mer. Our pseudoalignment criterion can be computed against all references at once by intersecting the non-empty color sets of all $k$-mers in a read.

We chose to implement our own $k$-mer index. The main design goal was that the index should be memory-efficient to build and use, because the size of the reference dataset can be large. To this end, we index the $k$-mer sets as a *succinct colored de Bruijn graph*. The nodes of the graph represent $k$-mers and the edges represent $(k+1)$-mers. The graph is encoded with a variant of the BOSS representation [3] and each node is linked to the corresponding color set with a separate coloring data structure which is unique to our implementation. Each query read is aligned as both the reverse complement string and the forward string, and we return the union of the pseudoalignments of both directions. Figure 1 illustrates the approach.

A speciality of our implementation is that the construction can be done almost entirely on disk, using only a minimal amount of RAM. This is made possible by designing the construction pipeline around two well-studied primitive operations: $k$-mer counting and disk-based sorting. The next section gives the technical details of the index and the construction pipeline.

#### 3 Implementation details

The reference sequences are modeled as strings from an alphabet $\Sigma$ of size $\sigma$ (for DNA, $\Sigma = \{A,C,G,T\}$ and $\sigma = 4$). Let us denote the set of references with $T_1, \ldots, T_m$. First, we build the BOSS data structure of the de Bruijn graph, implemented in terms of the generic Wheeler graph framework introduced by Gagie et al. [4].

Let $T = T_1\$T_2\$\cdots T_m\$$ be a dollar-separated concatenation of the reference sequences, where the dollar is a special symbol such that $\$ \notin \Sigma$. Let $f_\ell(x)$ be

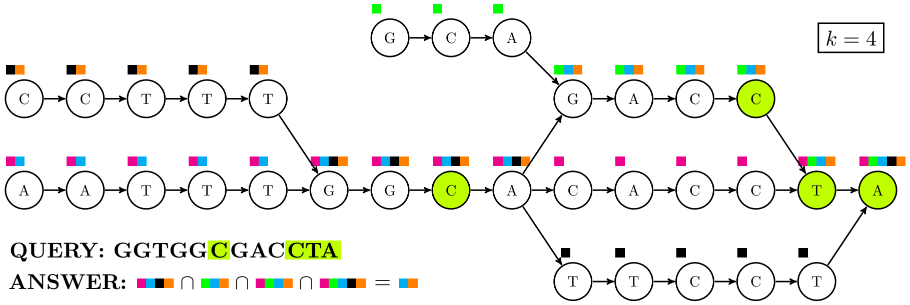


Figure 1: A colored de Bruijn graph of order $k = 4$. Each reference sequence in the graph is assigned a unique color. The color sets of nodes are drawn above the nodes. In the example, the query GGTGGCTGACCTA is pseudoaligned against the graph. Four of the $k$-mers of the query are found in the graph. The representative nodes of those $k$-mers are highlighted with green. The pseudoalignment returns the intersection of the color sets of the highlighted nodes.

the set of distinct characters that are found to the left of $k$-mer $x$ in $T$ and let $f_r(x)$ be the set of distinct characters that are found to the right of $x$ in $T$.

To build the Wheeler graph data structure, we iterate the sets $f_\ell(x)$ and $f_r(x)$ in colexicographic[1] order of the $k$-mers $x$ of $T$. To do this, we first list all distinct $(k+2)$-mers of $T$ to disk. Then we sort the $(k+2)$-mers $x$ in increasing order of the colexicographic rank of the middle $k$-mer $x[2..(k+1)]$ using a disk-based sorting algorithm. Next, we stream the sorted $(k+2)$-mers from disk. For every run of $(k+2)$-mers with an identical middle $k$-mer $y$, we collect the sets $f_\ell(y)$ and $f_r(y)$ by looking at the first and last characters of the $(k+2)$-mers in the run. Building the Wheeler graph data structure is straightforward from this information.

After this, we have a working index of the de Bruijn graph $(V, E)$ of the references $T_1, \ldots, T_m$. If $(u, v) \in E$, we call $u$ a predecessor of $v$, and $v$ a successor of $u$. Next, we add the colors to nodes of the graph. To eliminate redundancy, we only store colors for a subset $V' \subseteq V$, where $v \in V'$ iff at least one of the following conditions hold:

1. Node $v$ represents the first $k$-mer of a reference sequence.
2. A predecessor of $v$ represents the last $k$-mer of a reference sequence.
3. Node $v$ has multiple predecessors
4. Node $v$ has a predecessor that has multiple successors.

[1] The colexicographic order of strings is like the standard lexicographic order, but characters are compared starting starting from the end. The index can be build with either lexicographic or colexicographic sorting, but we choose to follow the colexicographic convention of the Wheeler graph framework. The indexed graph can be traversed in both directions.

If $v \notin V'$, then its color set has to be the same as its predecessor's color set. We can find out the color set of $v$ by walking backward to the nearest node $u \in V'$. Node $u$ is guaranteed to exist because the first node of every reference sequence is always in $V'$. The nodes in $V'$ can be found and marked by using the BOSS index.

However, with this setup, finding node $u$ might take a long time if we are in the middle of a long unitig (non-branching path), so we also store the color sets for some nodes inside long unitigs. Let $S$ be the set of nodes such that the distance backward to the nearest node in $V'$ is an integer multiple of $s$ for some global integer parameter $s$. We also store the color sets for all nodes in $S$. This way, we can find a color set of a node in at most $s$ backward steps. The sampling parameter $s$ can be tuned to obtain different time-space tradeoffs.

The color sets are computed with two disk-based sortings as follows. Assume we have marked all nodes in $V' \cup S$. Assign the reference sequences $T_1, \ldots, T_m$ colors such that the color of sequence $T_i$ is $i$. For each $i = 1, \ldots, m$, walk the de Bruijn graph according to $T_i$ using the constructed BOSS index, and for each node $v \in V' \cup S$ encountered, print to disk a pair $(v, i)$. After all sequences $T_i$ have been processed, sort the pairs on disk by the node identifiers $v$, and scan the sorted list, writing to another file pairs $(v, C_v)$, where $C_v$ is the list of colors of node $v$. Then, sort the new pairs by the color sets and scan the resulting sorted list to obtain a list of pairs $(X_v, C_v)$, where $X_v$ is the set of nodes with color set $C_v$. Finally, store all distinct color sets to a file, and for each node in the sets $X_v$, store a pointer to the corresponding color set.

It remains to be described how the color sets are stored in a succinct and accessible way. Let us denote the set of distinct color sets with $\mathcal{C} = \{C_1, \ldots, C_{|\mathcal{C}|}\}$. The color sets are stored in a concatenated form $C_1 \cdots C_{|C|}$. We mark with a bit vector all positions in the concatenation where a new color set starts, and index the bit vector for constant-time select queries to be able to locate the $i$-th distinct color set in constant time. A pointer to color set $C_i$ is just the integer $i$, which can be represented in $\lceil \log |\mathcal{C}| \rceil$ bits. By choosing the sampling parameter $s = \lceil \log |\mathcal{C}| \rceil$, the size of $S$ is at most $|V| / \log |\mathcal{C}|$, so the total size taken by the sampling pointers is only $|V|$ bits, and we obtain a worst-case color set lookup time of $O(\log s) = O(\log |\mathcal{C}|)$. With this, the whole coloring data structure takes on the order of $|V'| \log |\mathcal{C}| + |V| + \sum_{C \in \mathcal{C}} |C| \log m$ bits of space. The Wheeler graph data-structure takes $|V| \log \sigma + 2|V| + \sigma \log |V| + o(|V| \log \sigma)$ bits space, where $\sigma$ is the size of the alphabet.

Most of the heavy work is done by the subroutines for $k$-mer listing and for disk-based sorting. In our implementation, we used the highly optimized parallel tool KMC3 for $k$-mer listing, and a custom $\ell$-way disk-streaming mergesort with parallel merges for the sorting. The sorting implementation first divides the input into blocks that fit in the given RAM limit, sorts the blocks in RAM to disk, and then merges the blocks. Extra memory can speed up the sorting.

Any general purpose tool for the sorting and $k$-mer listing subroutines could be plugged into the pipeline with no changes to the rest of the pipeline. We believe this property could allow our construction pipeline to scale even to a distributed cluster of machines, as there are distributed implementations for both $k$-mer counting and sorting.

### 4 Performance

We benchmarked the construction performance of our implementation on a dataset of 3682 *Escherichia coli* genomes downloaded from the NCBI archives[2]. There were 19.0 billion nucleotides in this dataset.

Given 20GiB of RAM, Themisto builds the *E. coli* index for $k = 32$ in 6 hours and 16 minutes[3]. The main drawback is that the construction takes 375 GiB of disk space. Large disk usage is a common problem with sorting-based de Bruijn graph construction algorithms, such as the VARI-merge construction algorithm [5].

The final size of our index was 7.8GiB. The BOSS component of the index takes only 364MiB, and the rest of the space is taken by the coloring data structure. The concatenation of distinct color sets takes 6.6GiB of space. The distribution of the sizes of the color sets is shown in Figure 2. The index contains 325 million distinct $k$-mers.

Our implementation pseudoaligns reads from *E. coli* strains collected from across England [6] against the index at a rate of 1.4 billion nucleotides per hour using 8 threads, after loading the index into memory in 33 seconds. The alignment speed depends on the number matching $k$-mers and sizes of the color sets of the $k$-mers.

In comparison, Kallisto takes 4 hours and 57 minutes to construct an index for the same dataset, requiring as much as 287 GiB of memory. The index size on disk is 83 GiB, and 128 GiB in memory. The pseudoalignment throughput is approximately 2.1 billion nucleotides per hour using 8 threads, after loading the index to memory in 28 minutes. Table 1 summarizes key performance metrics for both Kallisto and Themisto on our benchmark.

| | Index in disk | Index in RAM | Indexing time | Indexing RAM | Indexing disk | Pseudoalignment throughput |
|---|---|---|---|---|---|---|
| Themisto | 7.8GiB | 7.8GiB | 6h 16min | 20GiB | 375GiB | $(1.4 \cdot 10^9)$ nt/h |
| Kallisto | 83GiB | 142GiB | 4h 57min | 287GiB | - | $(2.1 \cdot 10^9)$ nt/h |

Table 1: Themisto versus Kallisto on our benchmark dataset. The unit of throughput is nucleotides per hour.

[2]Assemblies from ftp://ftp.ncbi.nlm.nih.gov/genomes/genbank/bacteria/assembly_summary.txt with the organism name "*Escherichia coli*".

[3]Hardware: Intel Xeon E7-8890 CPU (2.2GHz, 60M Cache, 9.6GT/s QPI 24C/48T, HT, Turbo 165W) with 48 × 64GB LRDIMM memory (2400MT/s, Quad Rank, x4 Data Width), running on top of a distributed Lustre file system.

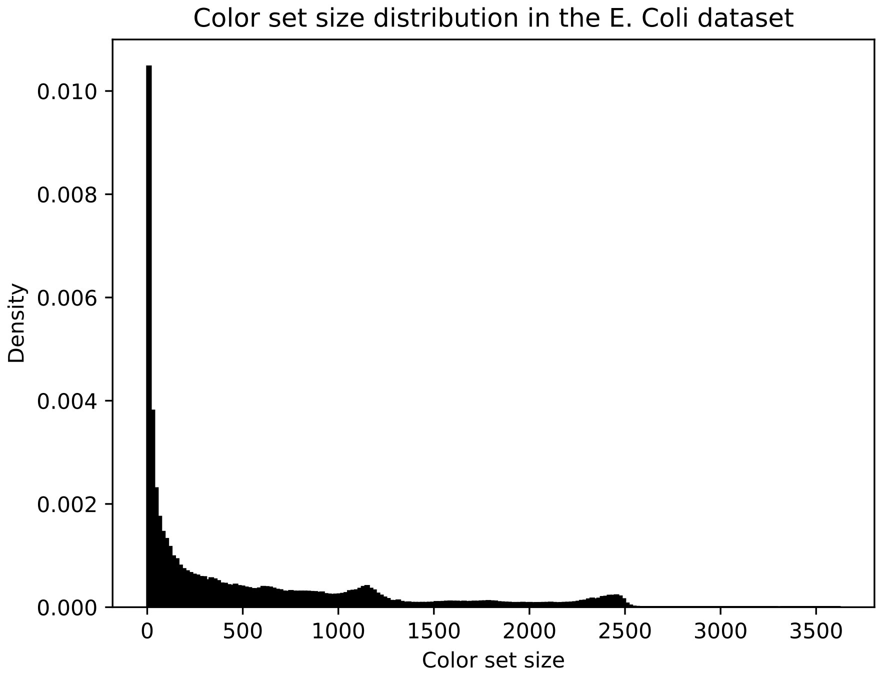


Figure 2: Color set size distribution for the dataset of 3682 E. Coli genomes each having a unique color.

[2] Camille Marchet, Christina Boucher, Simon J Puglisi, Paul Medvedev, Mikaël Salson, and Rayan Chikhi. Data structures based on k-mers for querying large collections of sequencing datasets. *bioRxiv*, page 866756, 2019.

[3] Alexander Bowe, Taku Onodera, Kunihiko Sadakane, and Tetsuo Shibuya. Succinct de Bruijn graphs. In *International workshop on algorithms in bioinformatics*, pages 225–235. Springer, 2012.

[4] Travis Gagie, Giovanni Manzini, and Jouni Sirén. Wheeler graphs: A framework for BWT-based data structures. *Theoretical computer science*, 698:67–78, 2017.

[5] Martin D Muggli, Bahar Alipanahi, and Christina Boucher. Building large updatable colored de Bruijn graphs via merging. *Bioinformatics*, 35(14):i51–i60, 2019.

[6] Teemu Kallonen, Hayley J Brodrick, Simon R Harris, Jukka Corander, Nicholas M Brown, Veronique Martin, Sharon J Peacock, and Julian Parkhill. Systematic longitudinal survey of invasive Escherichia coli in England demonstrates a stable population structure only transiently disturbed by the emergence of ST131. *Genome research*, 27(8):1437–1449, 2017.
