## Supplementary Figures 1-6 for "Genomic Epidemiology with Mixed Samples"

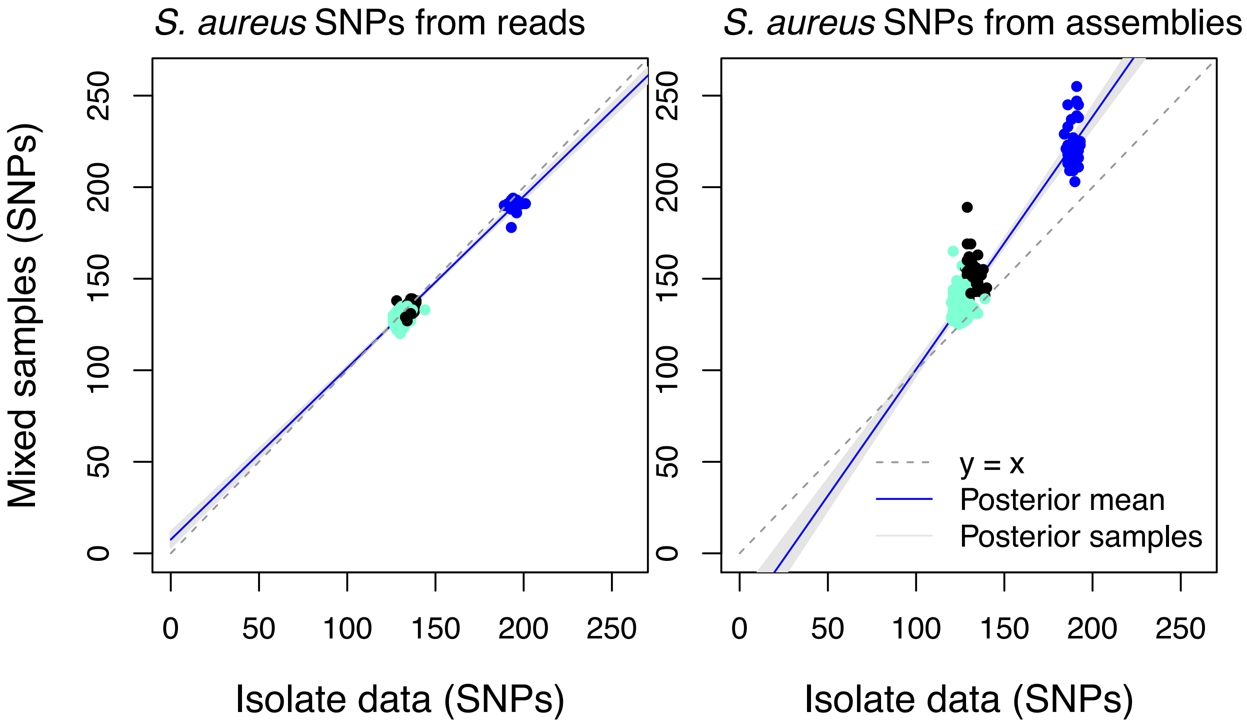


**Supplemental Figure 1** ***S. aureus*** **SNPs called from reads vs. assemblies obtained from the mGEMS pipeline.** SNPs in the panel on the right side were called from mixed samples processed with the standard mGEMS pipeline, while the panel on the left side omitted the final assembly step, which corresponds to the results presented panel c) of Figure 3 in the main manuscript. The colours indicate the different *Staphylococcus aureus* ST22 sublineages. The blue line is the posterior mean while the shaded area contains the 95% posterior credible region calculated from 10 000 posterior samples from a Bayesian regression model with the SNPs from the mGEMS-processed samples as the response and the SNPs from the isolate sequencing data as the sole explanatory variable.

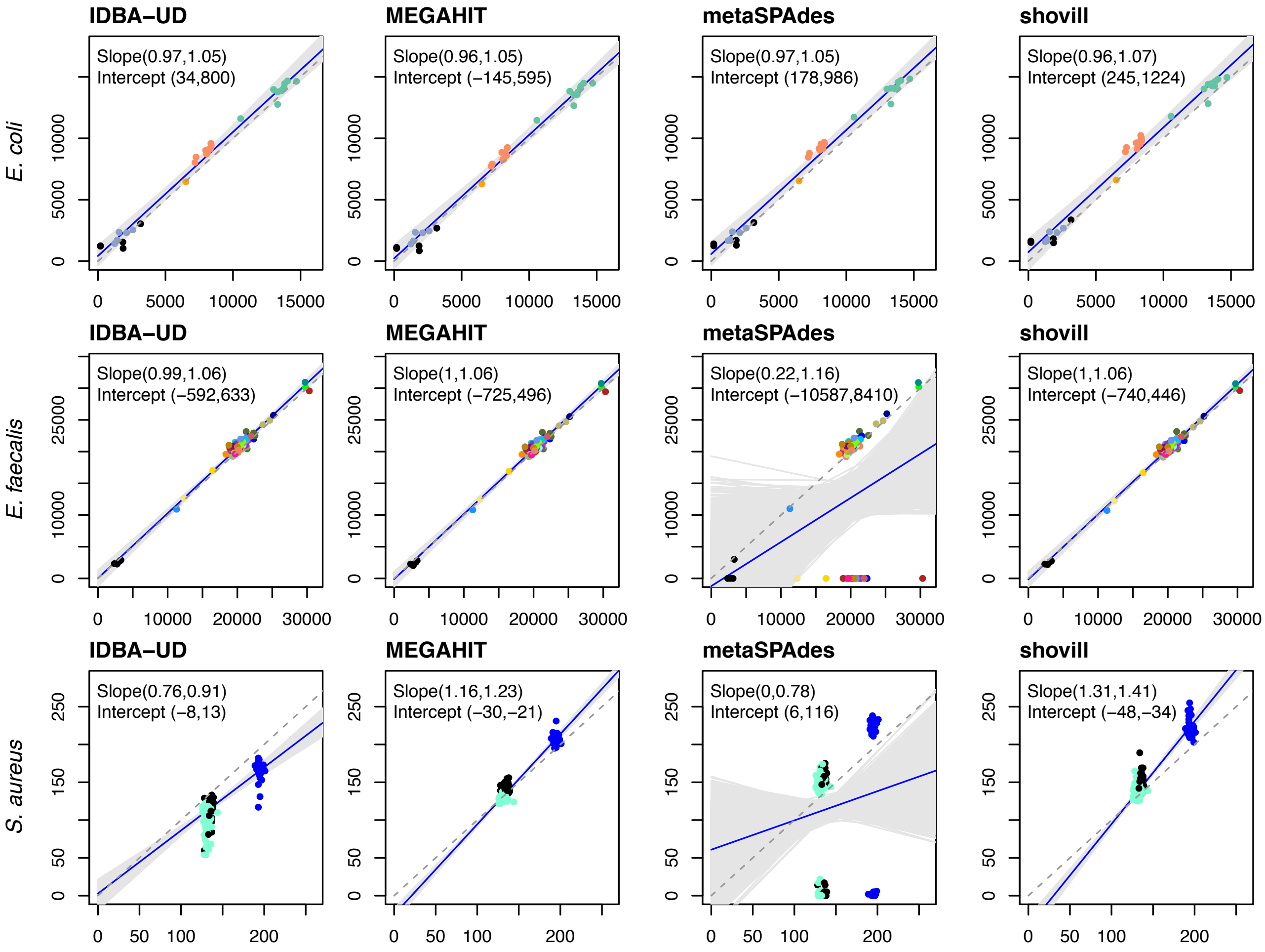


**Supplemental Figure 2 SNP calling with the mGEMS pipeline using different assemblers.** The plot shows the results of switching the Shovill assembler in the standard configuration of the mGEMS pipeline to a metagenomic assembler (IDBA-UD, MEGAHIT, or metaSPAdes, columns of the plot) and using the resulting assembly to call SNPs in the reference genome for each of the three species (*Escherichia coli, Enterococcus faecalis, and S. aureus;* rows of the plot). Expected results of the SNP calling (obtained from the isolate reads) are displayed on the horizontal axis. The different colours correspond to different lineages within the species. The blue line is the posterior mean while the shaded area contains the 95% posterior credible region calculated from 10 000 posterior samples from a Bayesian regression model with the SNPs from the mGEMS-processed samples as the response and the SNPs from the isolate sequencing data as the sole explanatory variable. The ‘Slope’ and ‘Intercept’ text in each subplot shows the 95% posterior credible region for both parameters as estimated by the Bayesian regression model.

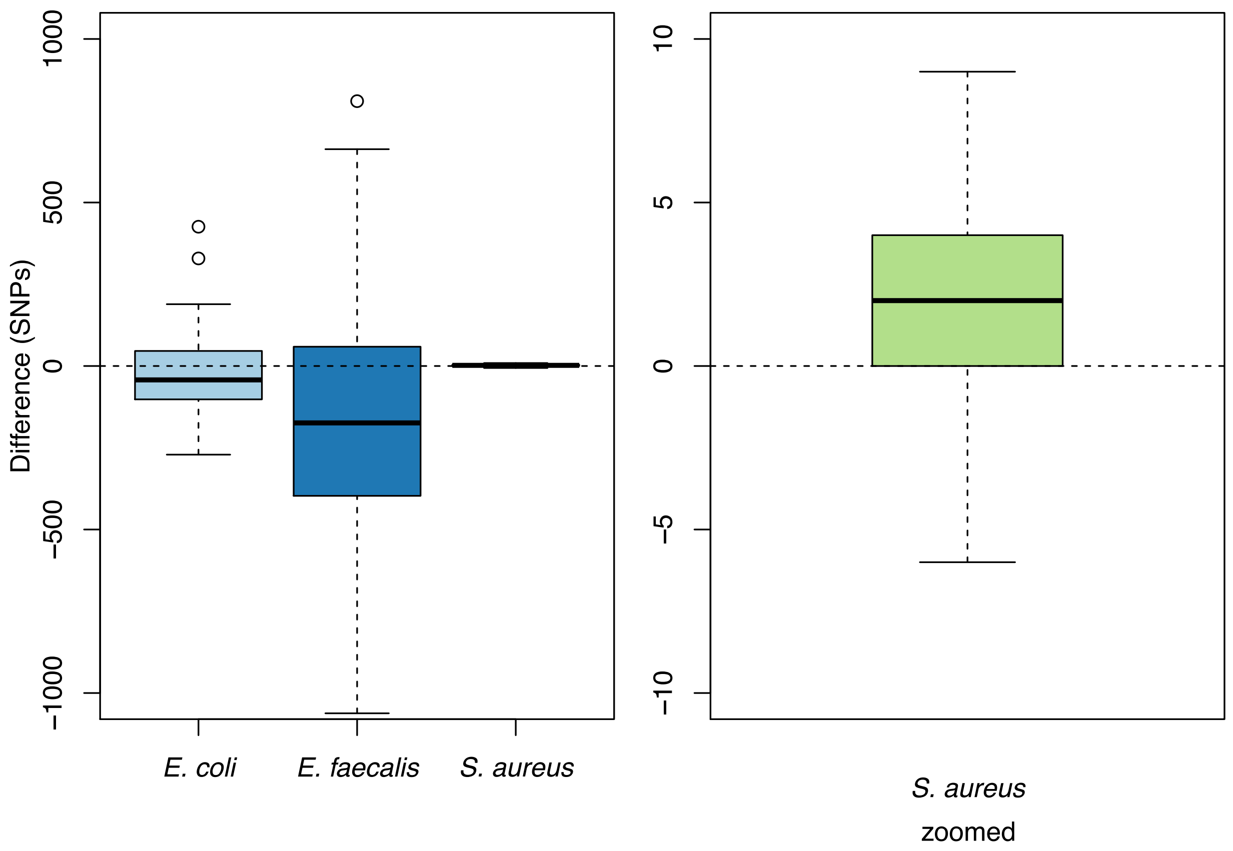


**Supplemental Figure 3 Difference in split-*k*-mer-SNPs called in the reference genome from isolate reads versus mixed samples binned with the mGEMS pipeline.** The panel on the left shows the results for all three species (*E. coli, E. faecalis,* and *S. aureus*), coloured by species. The panel on the right shows a zoomed-in view of the *S. aureus* results. The difference was calculated by subtracting the number of split-*k*-mer SNPs obtained from the mixed samples from the value obtained from the isolate reads.

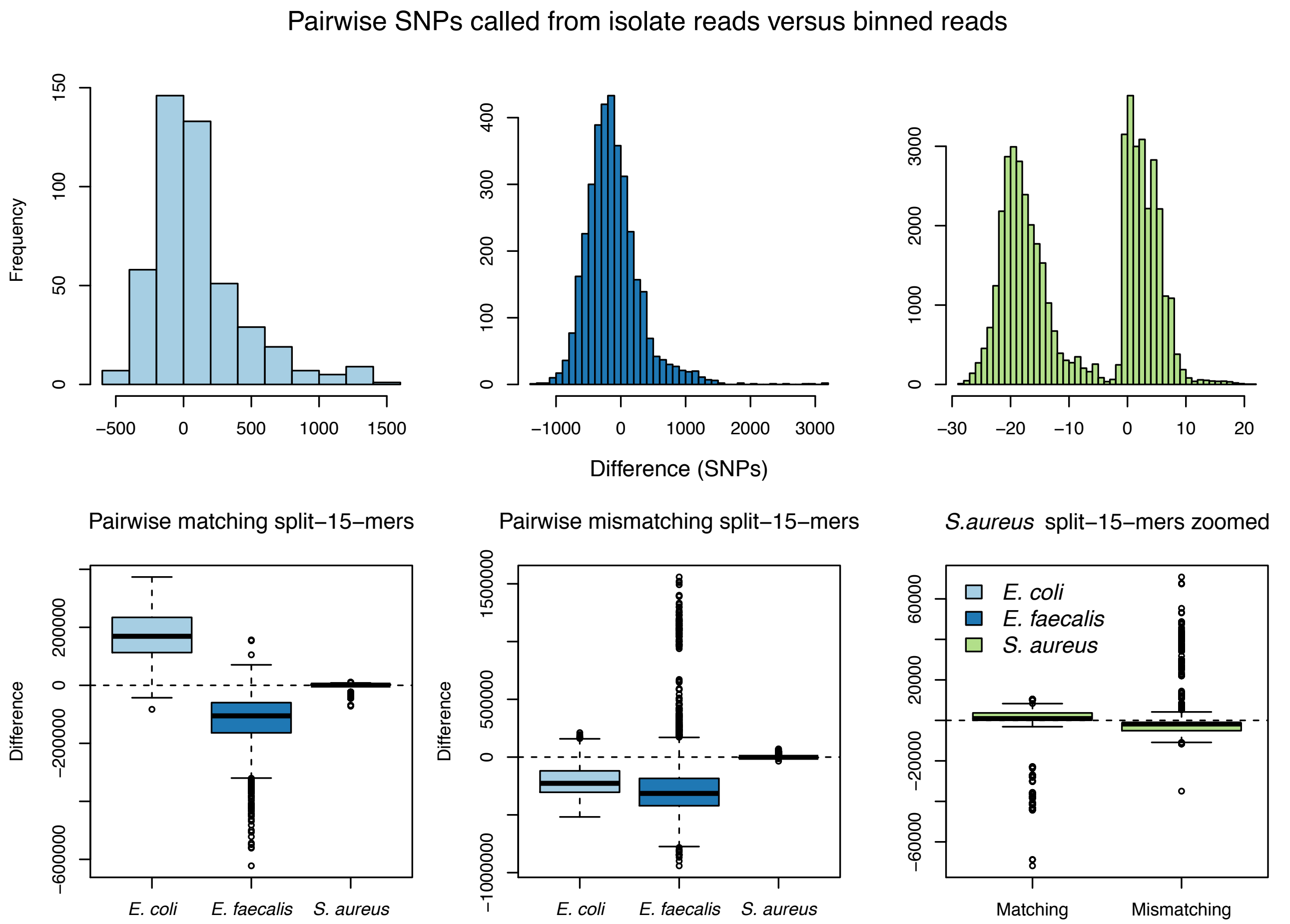


**Supplemental Figure 4 Pairwise SKA-SNPs called from isolate reads versus mixed samples binned with the mGEMS pipeline.** The top row shows the difference in the split-*15*-mer SNP counts calculated first pairwise within the set of isolate read samples, and then pairwise within the set of mixed samples processed with mGEMS. Similarly, the bottom row shows the differences in the numbers split-*15*-mers that are the same (matching) or different (mismatching) when the same pairwise analysis is performed. The values are coloured according to the species the samples originated from.

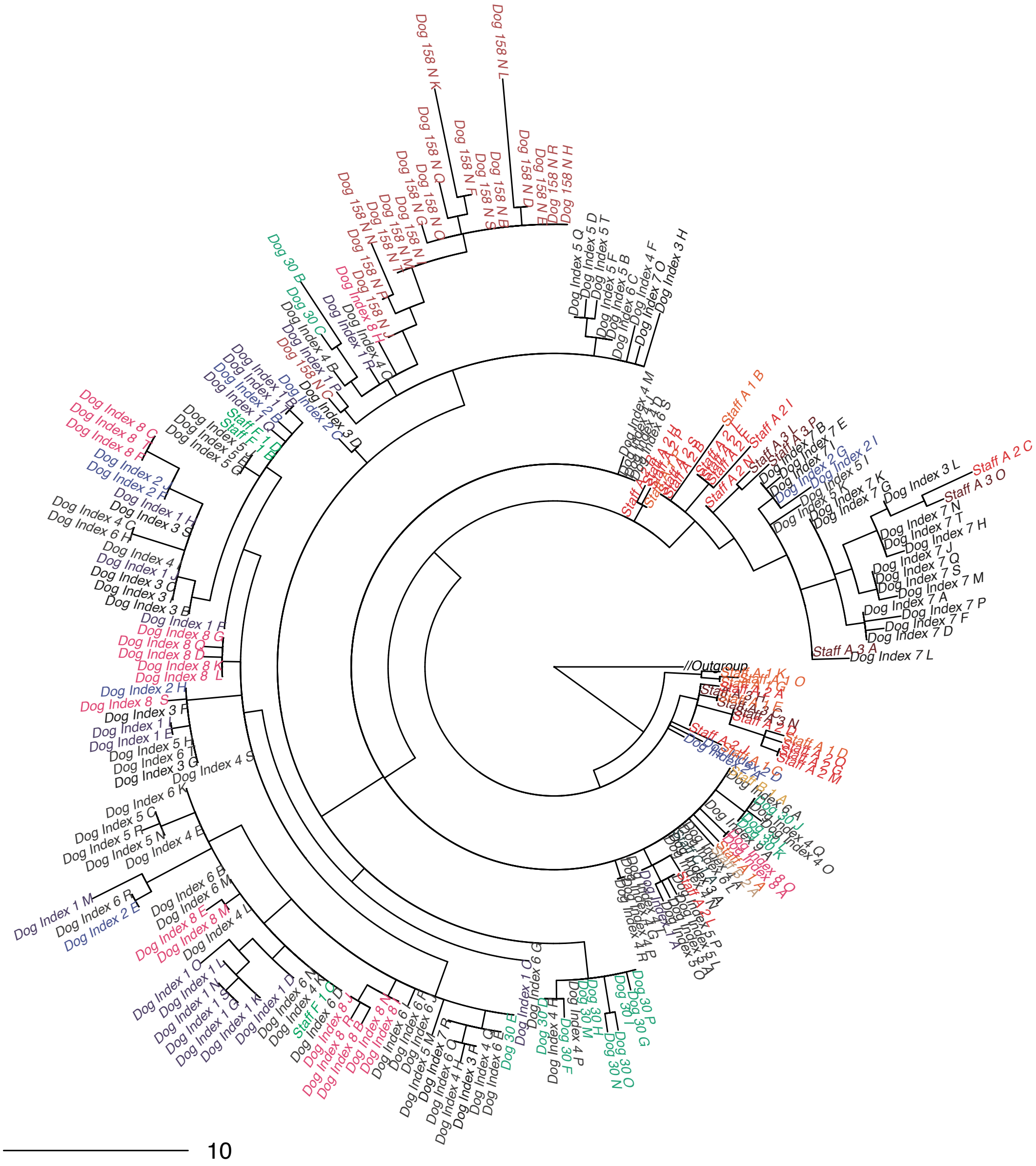


**Supplemental Figure 5 Midpoint-rooted maximum likelihood tree from core SNP alignment of *Staphylococcus aureus* ST22 isolate sequencing data showing the clade 1 strains.** Branches leading to clades 2 and 3, labelled *Outgroup*, were collapsed. Branch labels are coloured according to the plate the isolate sequencing data was originally picked from with darker shades indicating later sampling times. Branch lengths in the phylogeny scale with the mean number of SNPs obtained by multiplying the mean nucleotide substitutions per site on the respective branch (GTR+G4 model) with the total number of alignment sites. Leaves are labelled with the format: staff or patient, a letter indicating the donor, plate number (ascending in time), and a letter indicating the colony pick id.

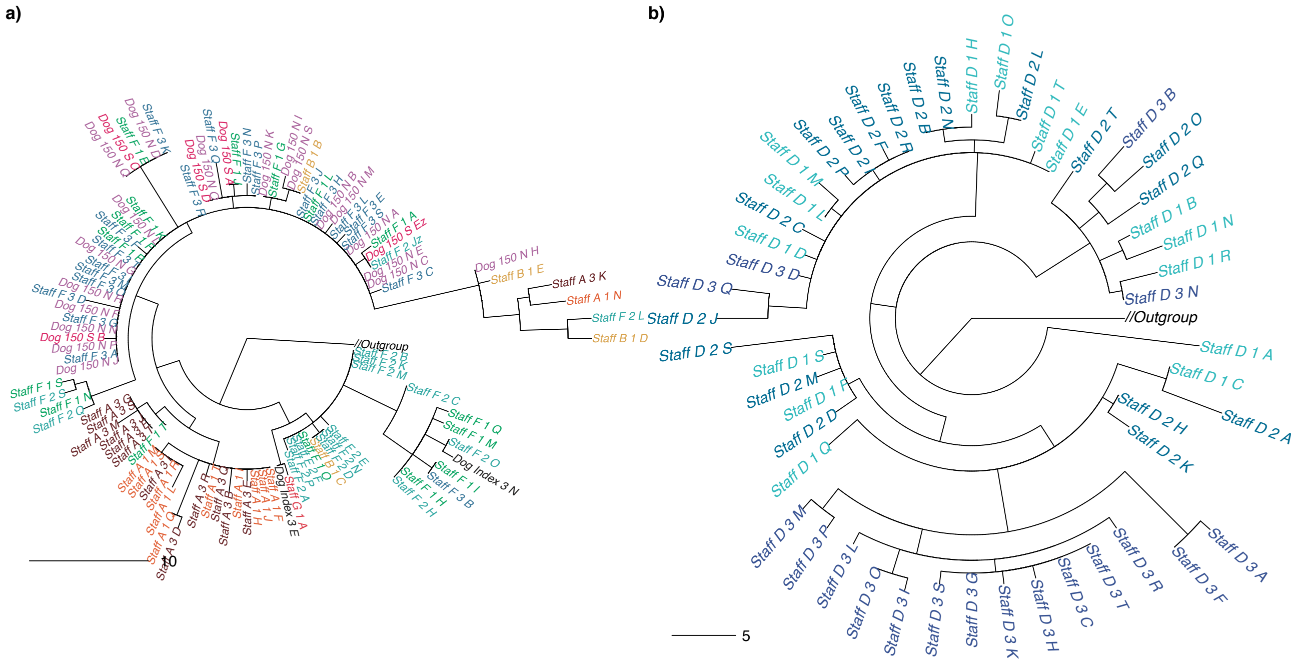


**Supplemental Figure 6 Midpoint-rooted maximum likelihood trees from core SNP alignment of *Staphylococcus aureus* ST22 isolate sequencing datashowing clade 2 and clade 3 strains.** Branches leading to clade 1 and clade 3 (panel a), or clade 1 and clade 2 (panel b), labelled Outgroup in both panels, were collapsed. Branch labels are coloured according to the plate the isolate sequencing data was originally picked from with darker shades indicating later sampling times. Branch lengths in the phylogeny scale with the mean number of SNPs obtained by multiplying the mean nucleotide substitutions per site on the respective branch (GTR+G4 model) with the total number of alignment sites. Leaves are labelled with the format: staff or patient, a letter indicating the donor, plate number (ascending in time), and a letter indicating the colony pick id.
